# Development of a hepatic cryoinjury model to study liver regeneration

**DOI:** 10.1101/2023.07.24.550437

**Authors:** Marcos Sande-Melon, David Bergemann, Miriam Fernández-Lajarín, Juan Manuel González-Rosa, Andrew G. Cox

**Affiliations:** Peter MacCallum Cancer Centre, Melbourne, Victoria, 3000, Australia; The Sir Peter MacCallum Department of Oncology, The University of Melbourne, Melbourne, Victoria, 3000, Australia; Cardiovascular Research Centre, Massachusetts General Hospital Research Institute, Charlestown Navy Yard Campus, 149, 13 Street, 02129 MA, USA; Harvard Medical School; Biology Department, Morrissey College of Arts and Sciences, Boston College, Chestnut Hill, MA 02467; Department of Biochemistry and Pharmacology, The University of Melbourne, Melbourne, Victoria, 3000, Australia

**Keywords:** Liver regeneration, cryoinjury, inflammation, proliferation, fibrosis, necrosis, apoptosis

## Abstract

The liver is a remarkable organ that can regenerate in response to injury. Depending on the extent of injury, the liver can undergo compensatory hyperplasia or fibrosis. Despite decades of research, the molecular mechanisms underlying these processes are poorly understood. Here, we developed a new model to study liver regeneration based on cryoinjury. To visualise liver regeneration at cellular resolution, we adapted the CUBIC tissue-clearing approach. Hepatic cryoinjury induced a localised necrotic and apoptotic lesion characterised by inflammation and infiltration of innate immune cells. Following this initial phase, we observed fibrosis, which resolved as regeneration re-established homeostasis in 30 days. Importantly, this approach enables the comparison of healthy and injured parenchyma with an individual animal, providing unique advantages to previous models. In summary, the hepatic cryoinjury model provides a fast and reproducible method for studying the cellular and molecular pathways underpinning fibrosis and liver regeneration.

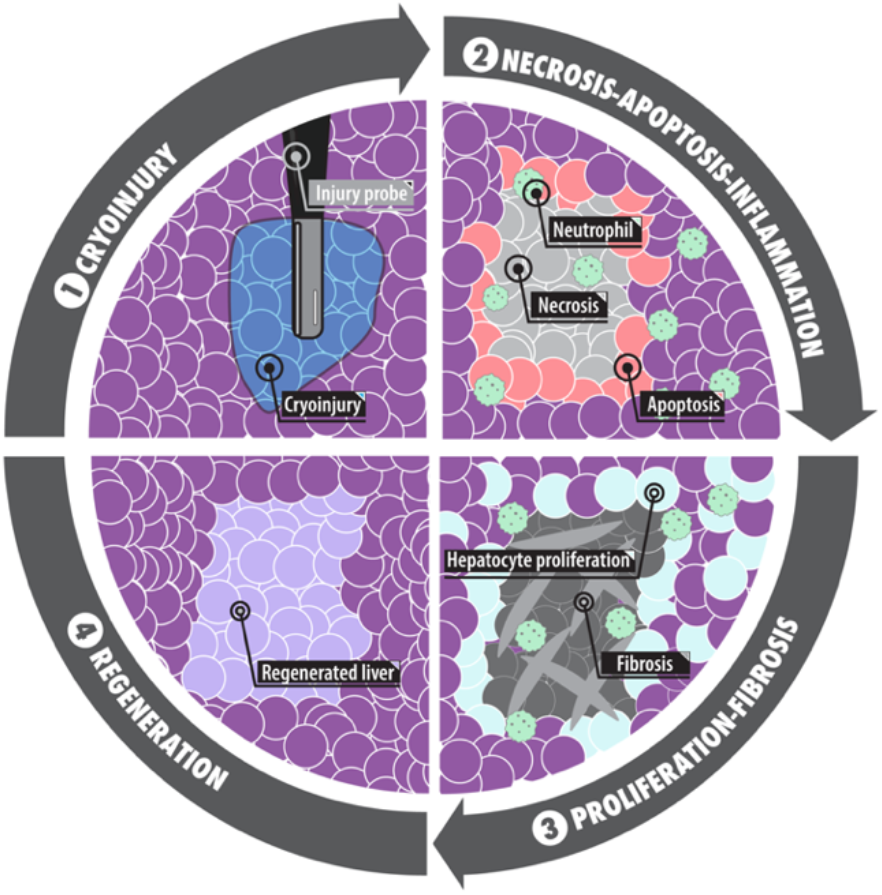

## Introduction

Regeneration is defined as the anatomical regrowth of an injured organ to restore homeostatic functions. Unlike most human organs, the liver exhibits an unparalleled capacity to regenerate. The process of liver regeneration is multifaceted, as it requires a complex tissue comprised of multiple cell types to sense the extent of the injury and mount an appropriate compensatory regrowth response^1^. Because of this complexity, the molecular underpinnings of liver regeneration are poorly understood.

Chronic liver diseases such as viral infection, alcohol, and non-alcoholic steatohepatitis often exhibit features of a maladaptive response to injury known as fibrosis ^2^. The phenomenon of fibrosis is a common wound-healing response in nature in which resident fibroblasts deposit extracellular matrix. In most cases, fibrosis inhibits the regenerative response of organs. However, zebrafish can resolve fibrotic scars to enable organ regeneration (e.g., cardiac regeneration following myocardial infarction^3–5^).

Several methods have been used to study liver regeneration in animal models. Among these methods, partial hepatectomy (PHx) is a surgical approach in which a liver lobe is resected, leaving the remnant liver lobes to regenerate^6–8^. In the PHx model of regeneration, inflammation or fibrosis has a limited role due to the absence of necrotic regions of tissue. The other dominant methods to study liver regeneration involve exposure to hepatotoxins that induce liver injury and, in some cases, fibrosis (e.g., CCl_49–11_, DMN^12–14^, TAA^15–17^, DDC^18^, APAP^19–22^ and ethanol^23–25^) ^26^. However, hepatotoxin-based approaches take months to develop liver injury and are therefore less amenable to rapid biological discovery. More recently, genetically encoded ablation models have been developed that enable specific liver cell types to be depleted (e.g., DTA^27^ and NTR ^28^). Each model has unique advantages and disadvantages; however, collectively they have led to remarkable advancements in our understanding of liver disease and regeneration.

Motivated to have a model of liver injury that is spatially localised yet retaining hallmarks of inflammation and fibrosis, we sought to adopt the cryoinjury approach successfully deployed in the cardiac field^3–5^. To this end, here we describe the development of a hepatic cryoinjury model of liver regeneration in this manuscript. We show that the hepatic cryoinjury model has no impact on survival and reproducibly leads to the liver regenerating within 30 days. Importantly, we observe that cryoinjury triggers an inflammatory phase associated with the clearance of necrotic and fibrotic tissue, followed by subsequent proliferation of hepatocytes at the injury site. A significant advantage of this model is that it enables the comparative analysis of healthy and injured parenchyma in a single animal. We anticipate that this hepatic cryoinjury model will facilitate the discovery of the cellular and molecular underpinnings of fibrosis and regeneration.

## RESULTS

### Hepatic cryoinjury in adult zebrafish induces localised cell death

We developed a novel method to study liver regeneration upon a localised cryoinjury of one of the liver lobes. Out of the three lobes present in the zebrafish liver, we chose the ventral lobe due to its surgical accessibility. In preparation for the surgery, zebrafish were immersed in anaesthetic and placed ventrally facing up in a foam holder (Figure 1A). To create a reproducible and stereotypical injury, we performed all injuries at the level of the anterior fins and toward the midline. First, a small incision was performed to expose the liver (Figure 1B). Once the liver was visible, the excess water was gently removed using a rolled tissue. Then, the cryoinjury probe, which was cooled in liquid nitrogen for one minute, was placed on the liver surface for 15 seconds until thawing (Figure 1C). Fish were quickly transferred into a freshwater tank and reanimated by pipetting water into their gills for 30 seconds (Figure 1D).

**Figure 1.**
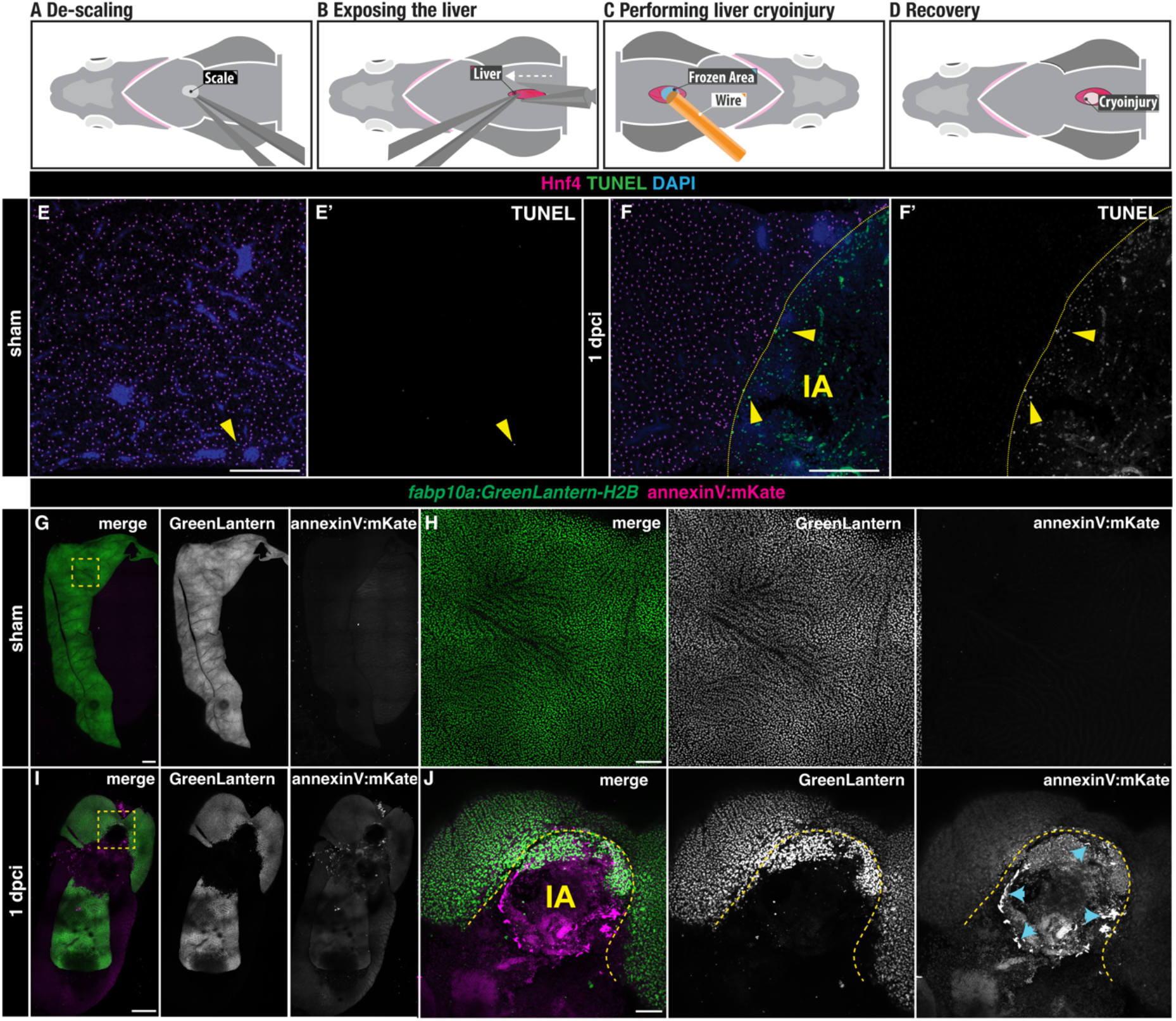
Liver cell death following cryoinjury. A-D) A simplified schematic illustrating the cryoinjury procedure in the zebrafish liver. A) The zebrafish liver is placed ventral side up to facilitate the surgery. B) A small incision near the midline exposes the ventral liver lobe. C) The frozen cryoprobe is applied to the liver surface for 15 seconds to induce injury. D) The damaged area in the liver appears as a blister-like structure at 1 dpci. E-F) TUNEL-staining of sham (E-E’) and injured (F-F’) liver sections at 1 dpci. IA, injured area. Yellow arrowheads: TUNEL^+^ cells; yellow dashed lines: border zone. G-J) *Tg(fabp10a: GreenLantern-H2B; annexinV:mKate) in toto* acquisitions of sham (G-H) and 1 dpci (I-J) livers. I-J) The IA (yellow dashed line) is identifiable by the absence of GreenLantern-H2B^+^ hepatocytes. Blue arrowheads: AnnexinV-mKate^+^ cells. Scale bars: 500 µm

To assess the extent of cauterisation following cryoinjury, we employed TUNEL staining in liver sections of adult zebrafish with hepatocytes labelled by Hnf4, comparing sham and 1 day post-cryoinjury (dpci, Figure 1E-F) livers. Sham-operated animals showed no TUNEL^+^ staining (Figure 1E-E’). In contrast, cryoinjury caused an injured area (IA) comprised of TUNEL^+^ cells surrounded by viable Hnf4^+^ hepatocytes (Figure 1F-F’). To determine the extent to which this injury model induces apoptosis, we performed cryoinjury in *Tg*(*fabp10a:H2B-GreenLantern; actb2:annexinV*-*mKate*) adult zebrafish in which apoptotic cells will be labelled with the mKate^+^ signal (Figure 1G-J). While we did not detect apoptosis in sham-operated animals (Figure 1G-H), AnnexinV-mKate^+^ apoptotic cells were observed around the injured area at 1 dpci, which can be easily detected by the localised loss of GreenLantern^+^ hepatocytes (Figure 1I-J). Interestingly, the annexinV-mKate^+^ cells were distributed in an annular field around the injured area but not in the central region (Figure 1J). Together, these results suggest that hepatic cryoinjury induces a localised cell death response characterized by a central necrotic core surrounded by a region of apoptotic cells at the injured border area.

### Liver regenerates upon cryoinjury in adult zebrafish

To understand the temporal dynamics of liver regeneration upon cryoinjury, we examined the extent of recovery at different times post-cryoinjury using *Tg*(*fabp10a: NLS-mCherry)* zebrafish livers (Figure 2A). Dissected livers were cleared, rendering them transparent, and subsequently scanned using *in toto* confocal imaging of the gastrointestinal block (Figure 2B-G, Supplementary Figure 1A-I, and Supplementary Figure 2A-I).

**Figure 2.**
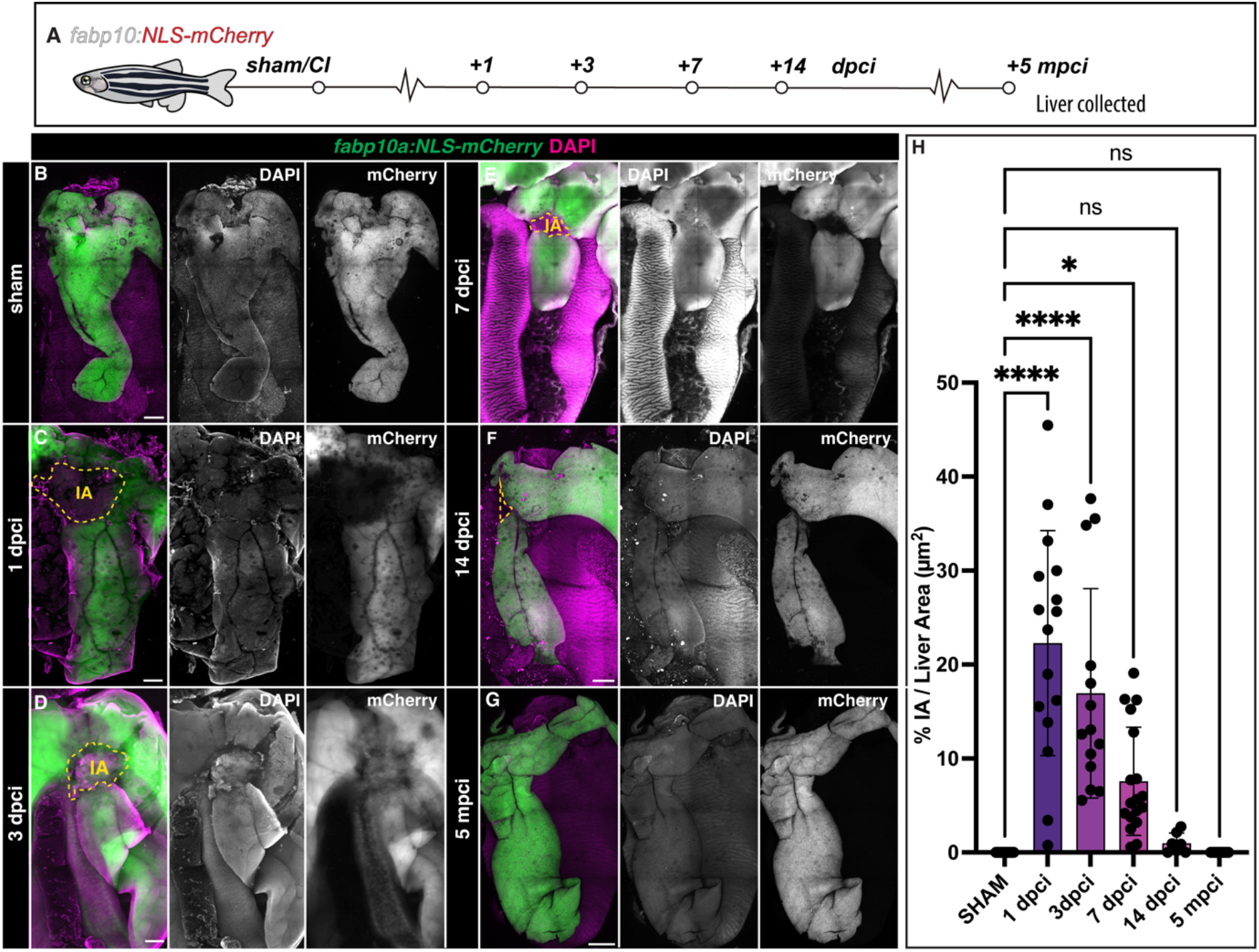
Progression of liver regeneration following cryoinjury. A) A simplified schematic illustrating the collection of livers following cryoinjury. B-G) Whole mount images of cleared zebrafish livers at the indicated stages of regeneration. Yellow dashed lines: border zone. H) Quantification of the IA area compared to the visible liver parenchyma area (n= 16, 16, 14, 18, 7, and 12 bars indicate mean, *p*-values: one-way ANOVA followed by Tukey’s multiple comparisons test). Scale bars: 500 µm.

Hepatic cryoinjury induced severe damage in the ventral lobe at 1 dpci, which was evident in histological sections and by the loss of the hepatocyte reporter fluorescence (Figure 2B-C, Supplementary Figure 2B-C, and Supplementary Figure 3C-E). Despite the extent of the injury, ∼93% of the animals survived this surgical procedure (Supplementary Figure 2O; n= 444). We observed a gradual recovery of mCherry^+^ hepatocyte expression from 3 dpci onwards, reducing the injured area’s size by confocal microscopy and histology (Figure 2D, Supplementary Figure 2D, and Supplementary Figure 3F).

At 7 dpci, when other liver injury models have fully regenerated^7,8,20,29^, the lesion is still clearly visible in cryoinjured livers (Figure 2E, Supplementary Figure 2E, Supplementary Figure 3H). The injured area is almost fully repaired at 14 dpci, as regeneration progressed (Figure 2F and Supplementary Figure 2F). Prolonged follow-up at 21 dpci (Supplementary Figure 2G), 3 months post-cryoinjury (mpci; Supplementary Figure 2H), 5 mpci (Figure 2G), 8 mpci (Supplementary Figure 3I), and 1 year post-cryoinjury (ypci; Supplementary Figure 2I) showed a fully regenerated liver parenchyma. Quantification of the injured area as a percentage of the total liver area at the different time points collected through this study indicates that the liver regenerates progressively over two weeks (Figure 2H).

To determine whether sex or age are biological variables that modify the response of the liver to cryoinjury, we next compared the injury area in male and female zebrafish cryoinjured at either 4 or 9 months of age (Supplementary Figure 2J). Although we found that injuries in females tended to be smaller than those of males at 7 dpci, we did not find significant differences among groups (Supplementary Figure 2J-M, and Supplementary Figure 2P).

Next, to ascertain whether pre-existing hepatocytes give rise to the regenerated hepatic parenchyma, we generated a hepatocyte-specific doxycycline-inducible Cre line for lineage tracing hepatocytes (Tg(*fabp10a:Tet-ON-Cre*)). We treated Tg(*fabp10a:Tet-ON-Cre*; *ubb*:Switch) animals with doxycycline, which induced genetic labelling of virtually all hepatocytes before injury (Supplementary Figure 2Q-R). At 7 dpci, we observed only recombined hepatocytes surrounding the injury area, indicating that pre-existent hepatocytes are responsible for the regeneration of the liver upon cryoinjury (Supplementary Figure 2S-T). Collectively, these results establish the hepatic cryoinjury model as a reproducible approach to induce a spatially localised injury that is repaired over several weeks.

### Hepatic cryoinjury induces a transient fibrotic response

Fibrosis is an evolutionarily conserved mechanism of liver repair that is beneficial when transient but can be maladaptive in chronic liver disease^2^. Given the extent of cell death observed following cryoinjury, we wanted to determine whether this injury model involved a fibrotic response. To this end, we used the Fuchsin Orange G (AFOG) staining^5,30^ to determine changes in ECM in response to cryoinjury. In the sham zebrafish liver, we mostly detected collagen surrounding the basement membrane of major blood vessels (Figure 3A). At 1 dpci, the insulted liver tissue started to develop collagen and fibrin deposition at the IA (Figure 3B), which gradually increased (Figure 3C-E), reaching a fibrotic peak at 5 dpci (Figure 3D, 3H, and Supplementary Figure 3A-B). Immunofluorescence staining with a collagen type I specific antibody revealed the presence of interstitial collagen depositions in the IA at 5 dpci (Figure 3H-H’’’), which is absent in sham-operated animals (Figure 3G-G’’’). During the following two weeks, fibrosis was gradually cleared and was almost undetectable at 30 dpci (Figure 3F), when the structure of the liver parenchyma was fully restored. Quantification of the collagen staining confirmed a significant increase in collagen deposition at 5 and 7 dpci compared to sham-operated, 1 and 30 dpci livers (Figure 3I). These results suggest that hepatic cryoinjury in the zebrafish leads to the formation of a transient fibrotic deposition, which is resolved within 30 days.

**Figure 3.**
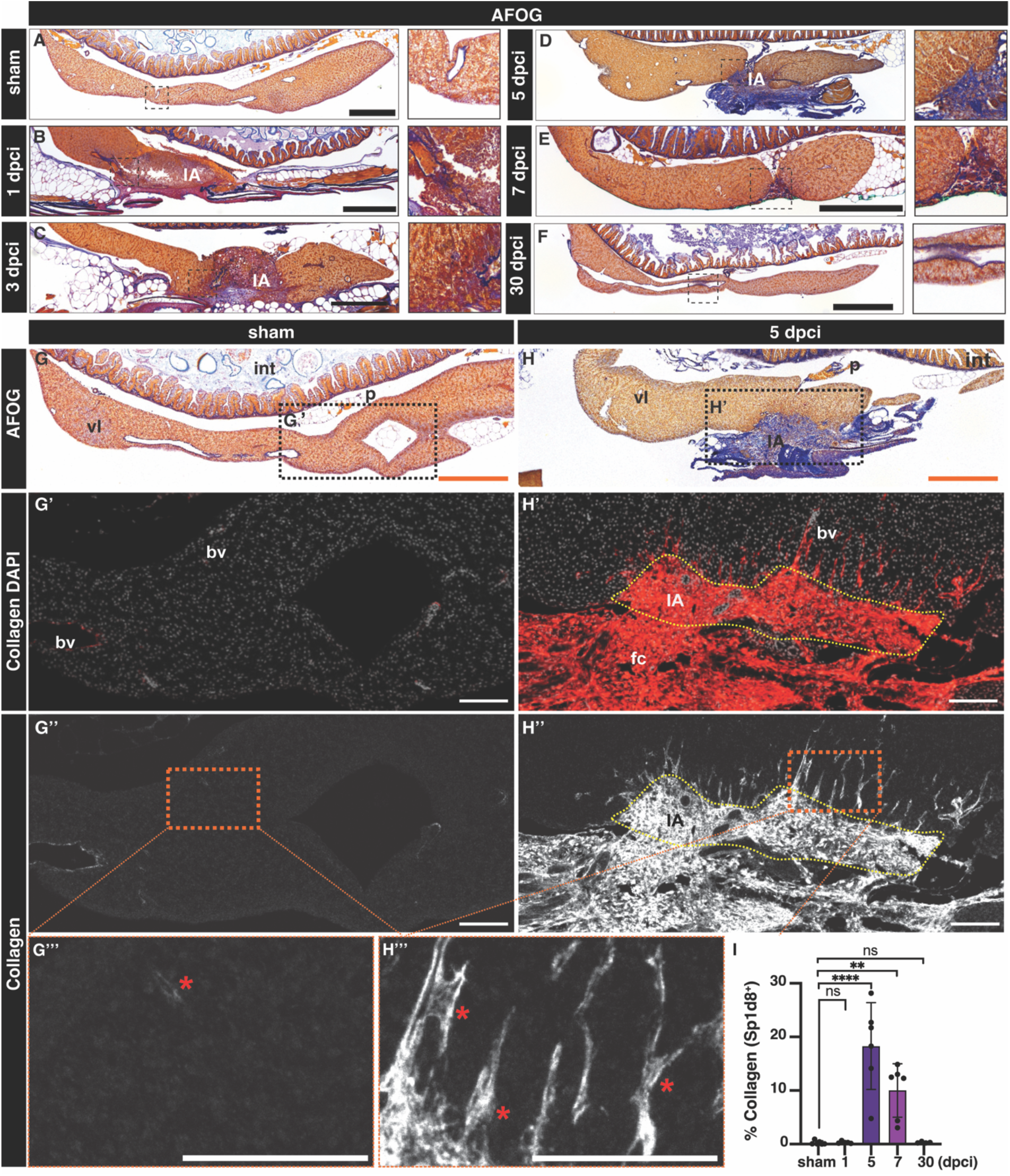
Transient fibrotic deposition during liver regeneration following cryoinjury. A-H) AFOG staining in sections of representative sham-operated (A,G) or injured livers at the indicated stages (B-H). Blue: collagen; Red: cell debris and fibrin. Anterior is towards the left, dorsal is towards the top. Details are shown at higher magnifications. G’H’) Adjacent sections from the samples showed in G and H, immunostained with an anti-Col1a1 antibody and counterstained with DAPI. Asterisks: Col1a1 deposition in the liver parenchyma. I) Quantification of the collagen area of livers from the indicated cohorts, normalized to the liver parenchyma area (n = 6, 5, 6, 6, and 4; error bars representing SD; *p*-values: one-way ANOVA followed by Tukey’s multiple comparisons test). Bv: blood vessel; fc, fibrotic cap; IA: injured area; int: intestine; p: pancreas; vl: ventral lobe. Scale bars: 100 (white), 500 µm (orange and black).

### Hepatic cryoinjury triggers a transient inflammatory response

The initial phase of liver regeneration is often associated with inflammation as leukocytes infiltrate the injured liver tissue^2,31^. Leukocytes are responsible for the phagocytosis of apoptotic and necrotic cells and the secretion of cytokines that promote hepatocyte survival and proliferation during liver tissue repair. To assess whether hepatic cryoinjury caused inflammation, we stained liver sham-operated and injured sections of *Tg*(*fabp10a*:NLS-mKate) zebrafish, carrying hepatocyte-specific mKate^+^ nuclear transgene expression in combination with the pan-leukocyte marker L-plastin (Lcp1) antibody (Figure 4A). In homeostatic livers, we observed a few Lcp1^+^ leukocytes in the liver tissue (Figure 4B). However, we detected a rapid infiltration of inflammatory cells after injury at 1 and 3 dpci, both in the healthy parenchyma and the cryoinjured area (Figure 4C-D). Immune infiltration was maintained for 7 days post-injury in the healthy liver parenchyma and the insulted area (Figure 4E-F). The presence of leukocytes gradually decreased at 18 dpci (Figure 4G), and reached basal levels at 30 dpci (Figure 4H), when the damaged area was no longer detected. Analysis and quantification of sham-operated and injured liver sections at different timepoints with Lcp1 staining showed a significant increase in the number of infiltrated leukocytes at 1, 3, 5, and 7 dpci (Figure 4I).

**Figure 4.**
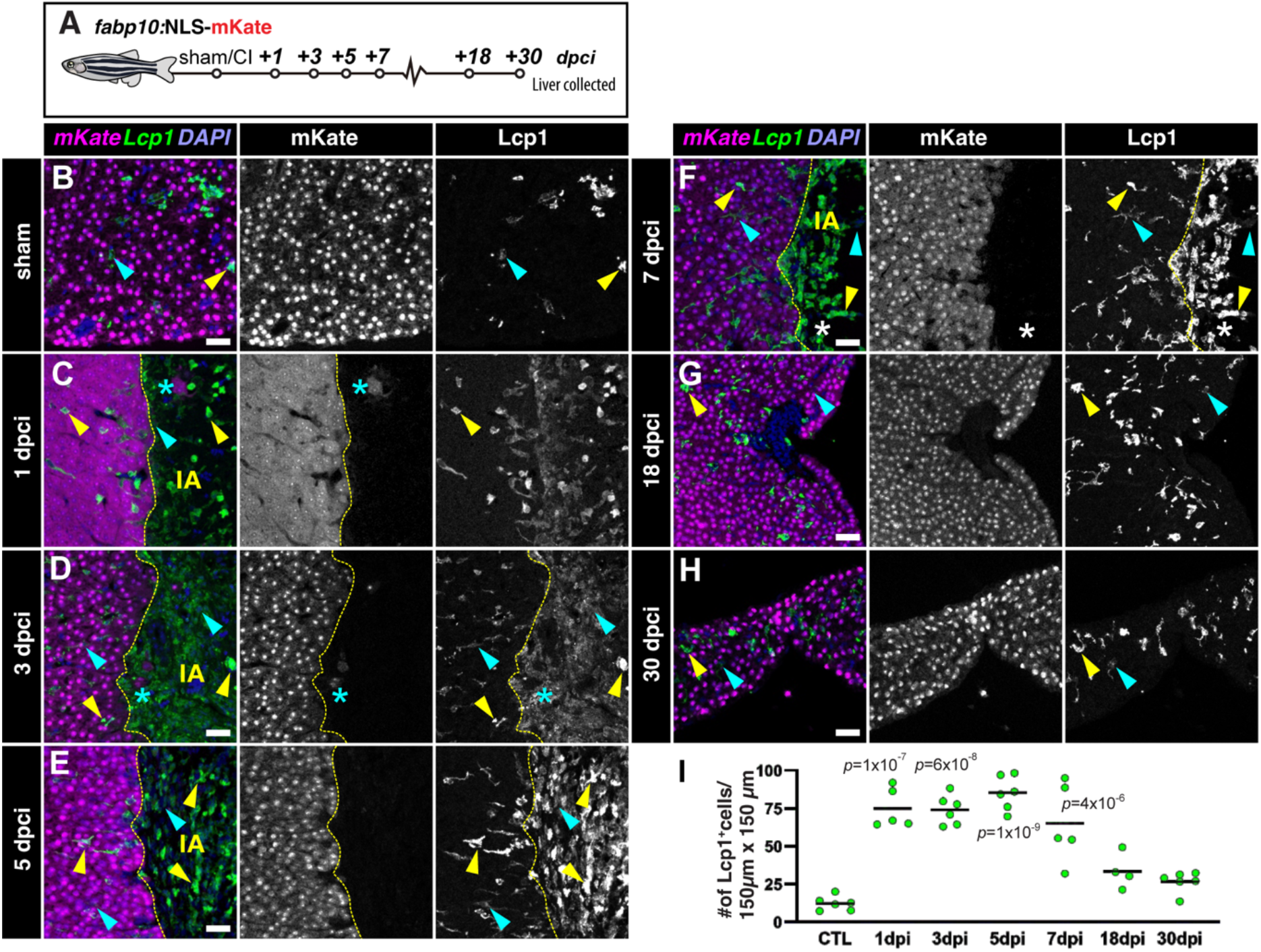
Cryoinjury induces the local and transient infiltration of leukocytes. A) Schematic representation of the experiment workflow. B-H) Sections of livers from *Tg(fabp10a:NLS-mKate)* animals at the indicated stages, immunostained to detect hepatocyte nuclei (mKate) and leukocytes (Lcp1) Cyan arrowheads: low signal Lcp1^+^cells; dashed yellow line: border zone; IA: injured area; yellow arrowheads: high signal Lcp1^+^ cells; Asterisks: spared hepatocytes surrounded by injured/necrotic tissue area. (I) Quantification of the number of Lcp1^+^ cells in designated regions (n = 6, 5, 6, 6, 5, 4, and 6; solid black line indicates the mean). *p*-values: one-way ANOVA followed by Tukey’s multiple comparisons test. Scale bars: 25 µm.

Using Lcp1, we gained some insights into the dynamics of the inflammatory response, but we lacked granular information about the specific populations and their potential role during regeneration. To characterize this process further, we stained sections from sham-operated and injured livers with antibodies that recognize Mpx, a neutrophil-specific marker (Supplementary Figure 4A). While we did not find neutrophils in homeostatic livers (Supplementary Figure 4B-B’), we detected infiltrated neutrophils at 1, 3, and 5 dpci specifically in the cryoinjured area but not in the contralateral lobe (Supplementary Figure 4C-E’). This response was transient, as analysis of livers at 7 dpci evidenced a reduction to basal levels by that stage (Supplementary Figure 4F-F’).

Macrophages are essential for regeneration of the zebrafish heart, limb, and brain^32-34^. To study whether macrophages are necessary for liver regeneration upon cryoinjury, we performed intraperitoneal injections (IP) of either PBS or clodronate liposomes to deplete macrophages during the regenerative process. Adult livers were collected to assess the injured area and were immunostained to detect Mfap4^+^ macrophages at 3 and 7 dpi (Supplementary Figure 5A). We did not find significant differences in the regenerative response in livers treated with clodronate liposomes (Supplementary Figure 5B-C, and 5J), albeit a noticeable reduction in Mfap4^+^ cells in the treated group (Supplementary Figure 5F-I). Collectively, these experiments confirm that hepatic cryoinjury is associated with the local and transient infiltration of leukocytes into the injured area.

### Cryoinjury induces both localised and distal compensatory hyperplasia

Two regenerative mechanisms could be at play in the cryoinjury model, namely compensatory hyperplasia, in which hepatocyte proliferation occurs throughout the parenchyma until liver mass reaches homeostasis, or epimorphic regeneration, in which hepatocytes proliferate in a spatially localised region around the injury. To understand the regenerative response upon hepatic cryoinjury, we examined hepatocyte proliferation. To identify proliferative hepatocytes, we stained sections of *Tg(fabp10a:NLS-mKate)* with the proliferating cell nuclear antigen antibody (PCNA; Figure 5A). Additionally, we performed BrdU pulse-labelling studies, injecting BrdU in sham and injured zebrafish at 2 dpci (Supplementary Figure 6A). Quantification of hepatocyte proliferation (mKate^+^/PCNA^+^) confirmed that there were few mKate^+^/PCNA^+^ cells (Figure 5B and 5I) and mCherry^+^/BrdU^+^ HCs (Supplementary Figure 6B,D and E) in sham-operated animals, consistent with the slow turnover of hepatocytes in adulthood during homeostasis^32–34^. In contrast, proliferating hepatocytes were significantly more abundant around the injured area during the first week of recovery after cryoinjury (Figure 5C-F and Supplementary Figure 6C-K).

**Figure 5.**
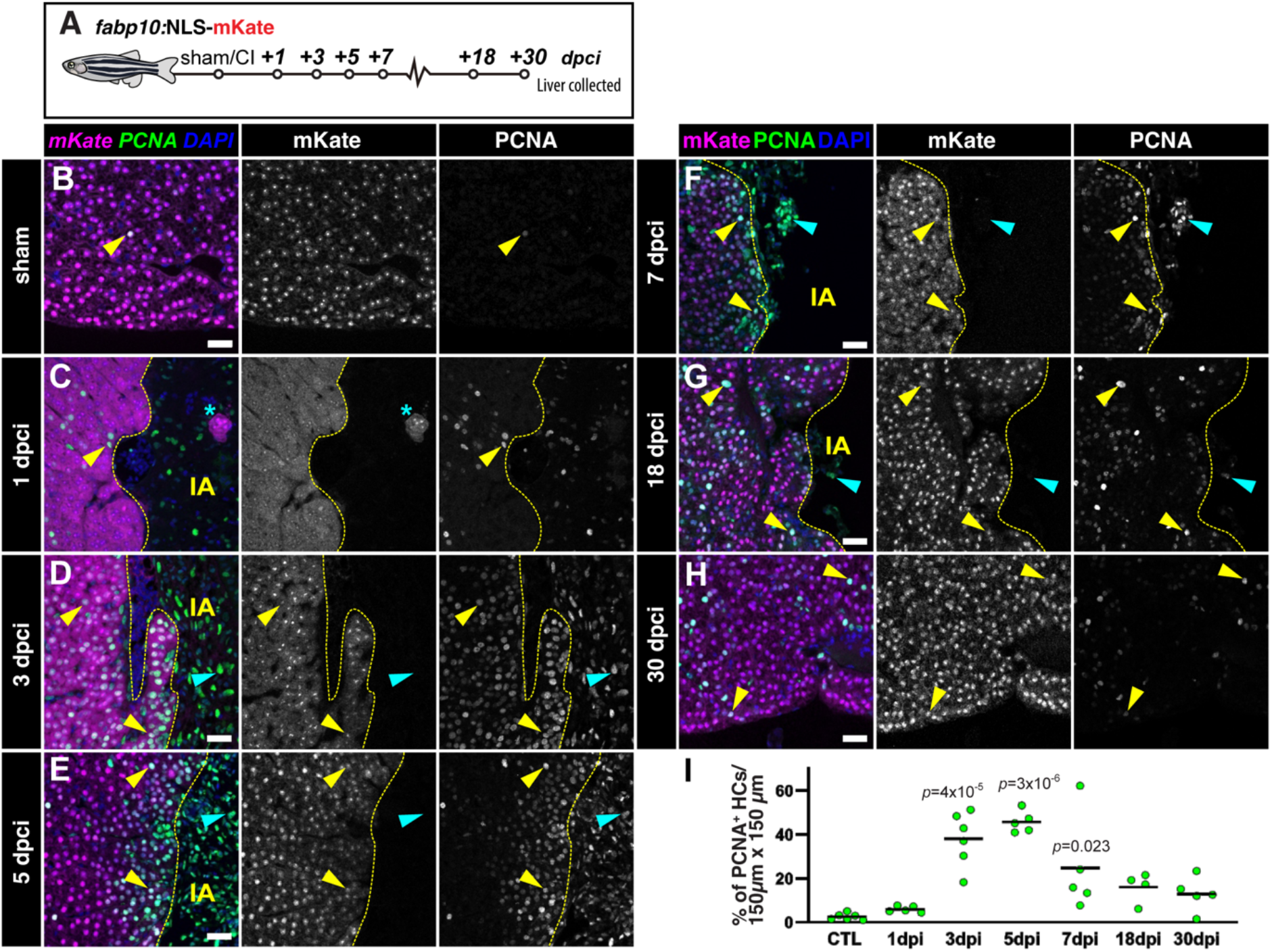
Local hepatocyte hyperplasia upon cryoinjury. A) Schematic representation of the experiment workflow. B-H) Liver sections from *Tg(fabp10a:NLS-mKate)* animals at the indicated stages, immunostained to detect proliferation (PCNA) and hepatocyte nuclei (mKate). Yellow arrowheads: proliferating hepatocytes. Cyan arrowheads: other cell types actively cycling. Dashed yellow line: separation between healthy and injured liver parenchyma; IA, injured area; Asterisks: spared hepatocytes surrounded by injured/necrotic tissue area. I) Hepatocyte proliferation index in the border zone at the indicated stages (n = 6, 5, 6, 5, 5, 4, and 5; solid black line indicates the mean). *p*-values: one-way ANOVA followed by Tukey’s multiple comparisons test. Scale bars: 25 µm.

Hepatocyte proliferation peaked between 3 and 7 dpci, which correlates with the timing of reduction in the IA (Figure 5I). Analysis of proliferating hepatocytes shows a localised hyperplasia, with a significantly higher density of cycling cells in areas surrounding the injured area compared to the contralateral lobes (Supplementary Figure 6D-F and 6I-K). This recovery of hepatocytes around the cryoinjured area is consistent with an epimorphic regeneration. Notably, we detected proliferating hepatocytes in the contralateral lobe (Supplementary Figure 6C-F and 6G-K), indicative of an adaptive compensatory hyperplasia. Hepatocyte proliferation decreased during later stages of regeneration to a baseline at 30 dpci (Figure 5G-H). Therefore, our data suggests that hepatic cryoinjury stimulates features of both epimorphic and compensatory hyperplasia during liver regeneration.

### The biliary and endothelial network is re-established during regeneration

Adaptive liver regeneration that recovers hepatic function requires appropriate restoration of the biliary and vascular network, which are comprised of biliary epithelial cells (BECs) and endothelial cells (ECs), respectively. To examine whether BECs regenerate following hepatic cryoinjury, we collected *Tg(fabp10a: H2B-GreenLantern)* livers at 1, 3, 7, and 21 dpci (Figure 6 A-F and Supplementary Figure 7 A-C), performed CUBIC clearing, and immunodetected the BEC marker Annexin A4 (Anxa4). Anxa4^+^ BECs were detectable throughout the liver parenchyma in sham livers (Figure 6A-A’’).

**Figure 6.**
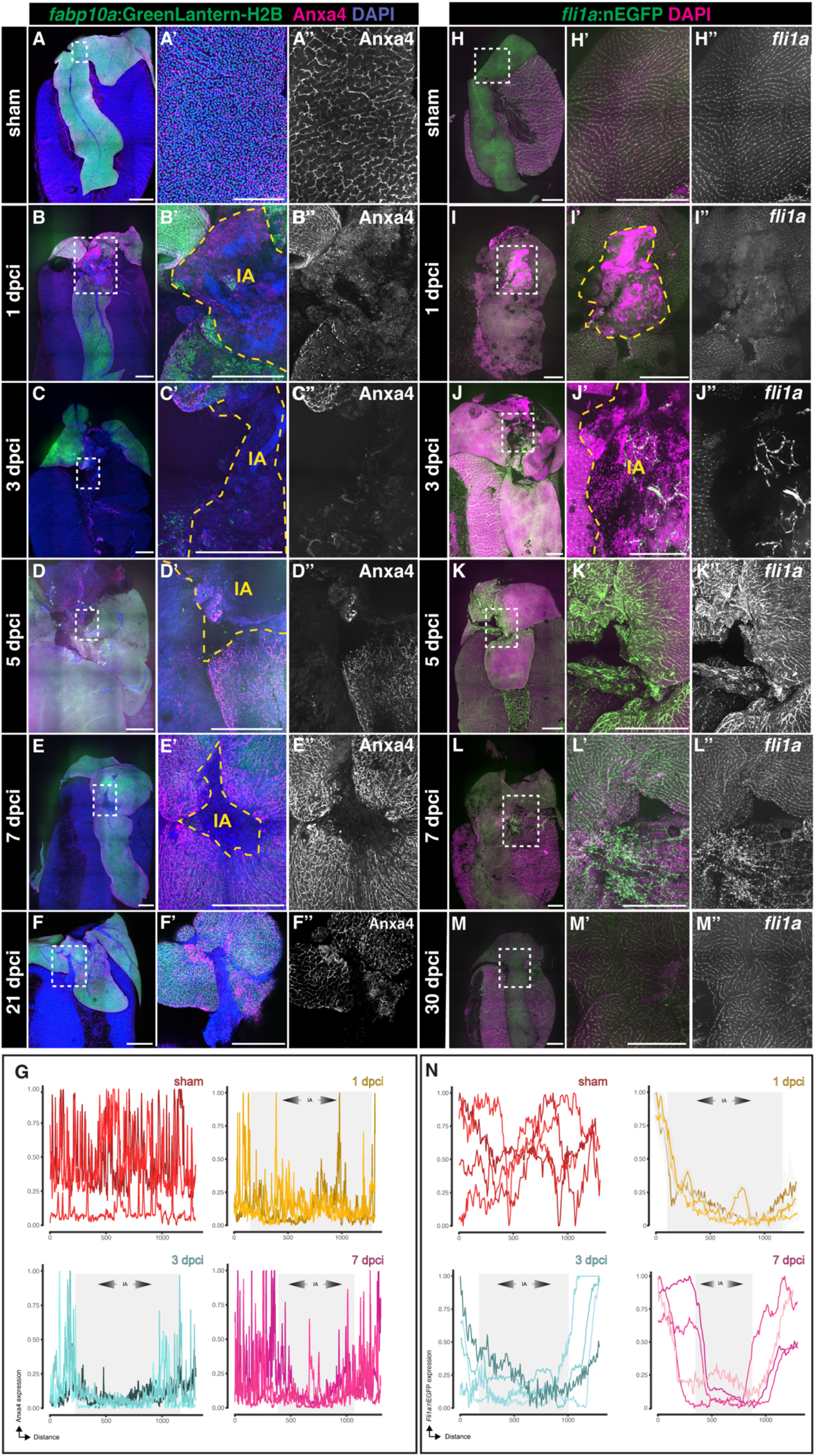
Biliary Epithelial Cells (BECs) and endothelial cells (ECs) recover upon cryoinjury. A-F) Whole mount imaging of *GreenLantern*^+^ hepatocytes and Anxa4^+^ biliary epithelial cells from sham (A) or injured (B-F) livers at the indicated stages. Details from boxed regions are shown magnified. G) Mean intensity profile of Anxa4 within the IA at the designated stages is represented by individual lines for each sample, with the IA delineated in grey (n= 4, 4, 4, 4). H-M) Whole mount acquisitions of GFP^+^ ECs from sham (H) or injured (I-M) livers at the indicated stages. Details from boxed regions are shown magnified acquisitions. N) Mean intensity profile of *fli1a*:nGFP within the IA at the designated stages is represented by individual lines for each sample, with the IA delineated in grey (n= 4, 4, 4, 4). Dashed line: border zone of the injured area, IA: injured area. Scale bars: 500 µm.

However, the distribution of Anxa4^+^ BECs was disrupted in the IA at 1 and 3 dpci, indicating that bile duct structures were affected during this procedure (Figure 6B-C, Figure 6G, and Supplementary Figure 7C). A progressive recovery of the biliary network was observed at 7 and 21 dpci, as the insult area gradually reduced in size (Figure 6E-F and Supplementary Figure 7C). Importantly, no differences were observed in the architecture of the biliary network in regenerated livers at 21 dpci compared to sham-operated animals (Figure 6A’’, Figure 6F’’), consistent with the regenerative response in hepatocytes. We documented a dynamic and significant increase in the proliferation of BECs after injury, peaking at 3 dpci (Supplementary Figure 6L-O). We also detected a modest but significant increase in BEC proliferation in the contralateral lobe at 3 dpci (Supplementary Figure 6P), suggesting that the regeneration of the biliary tree is predominantly driven by localized hyperplasia and not by compensatory growth.

To determine the dynamics of regeneration of the endothelial cells (ECs) after injury, we collected *Tg(fli1a:NLS-eGFP)* livers at 1, 3, 7, and 30 dpci (Figure 6F-J and Supplementary Figure 6 F-G). eGFP^+^ ECs were observed equally through the sham liver parenchyma (Figure 6F-F’’ and Supplementary Figure 6F-G). However, the distribution of ECs was affected in the IA at 1 dpci (Figure 6G-G’’ and Supplementary Figure 6F-G). Vascular recovery starts to be observed at 3 dpci, with branching morphogenesis of eGFP^+^ ECs appearing within the IA (Figure 6H-H’’ and Supplementary Figure 6F-G) at early stages, consistent with previous reports^35^. Interestingly, eGFP^+^ ECs are detected throughout the entire IA at 7dpci, revealing that vasculature regeneration precedes HCs and BECs regeneration (Figure 6I-I’’ and Supplementary Figure 6F-G). However, the endothelium appeared significantly disorganized in the IA compared to sham-operated animals (Figure 6I’’, Figure 6F’’), and this abnormal distribution persisted at 30 dpci (Figure 6J-J’’), even when the liver parenchyma was fully recovered. These findings demonstrate that hepatic cryoinjury disrupts the biliary and vasculature network, which is subsequently re-established during liver regeneration.

### Characterization of the transcriptional landscape following cryoinjury

To study the transcriptional changes that occur during liver regeneration, we performed bulk RNA-seq on the injured region in the ventral love and contralateral lobes of livers dissected from *Tg(fabp10a:NLS-mCherry)* animals following sham-operation, 1, 3, and 7 dpci (Figure 7A and Supplementary Figure 8A-B). At 1 dpci, we detected 438 upregulated differentially expressed genes (DEGs) and 981 downregulated genes when compared to sham-control liver (Figure 7B and 7E-F).

**Figure 7.**
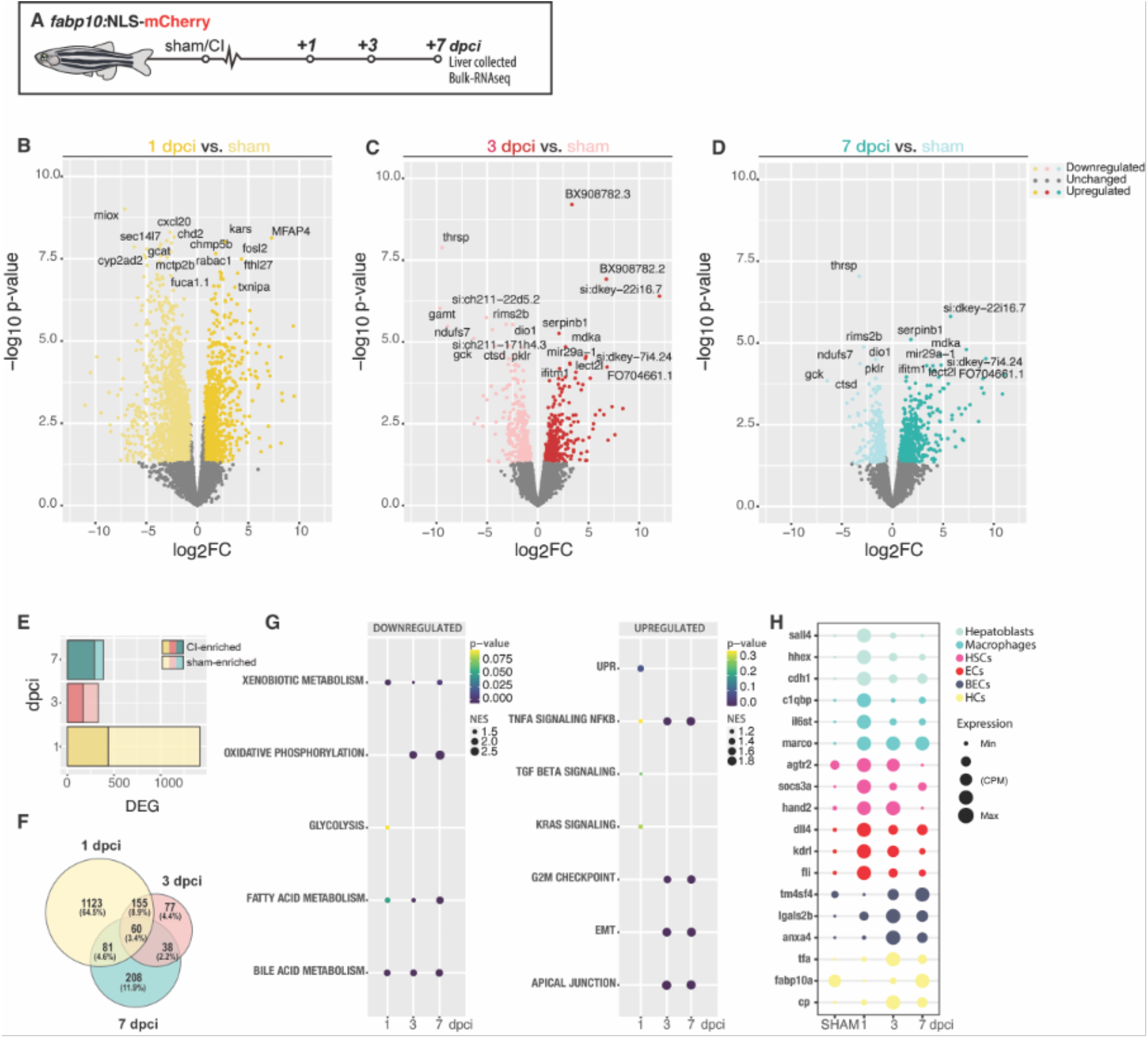
Transcriptional signatures of liver regeneration upon cryoinjury. A) Schematic representation of the experiment workflow. B-D) Volcano plots representing the comparison of 1, 3 and 7 dpci with sham adult zebrafish. DEGs (FC ≥1.5 (darker dots) or ≤-1.5 (lighter dots;); *P*≤0.05; with top DEG annotated). E) Bar plot representing the number of upregulated and downregulated DEG at 1, 3, and 7 dpci. F) Venn diagram representing DEG at 1,3, and 7 dpci. G) GSEA of liver cell types during liver regeneration. H) Dotplot representing the expression of key genes for specific liver cell populations upon cryoinjury.

Gene Set Enrichment analysis (GSEA) identified the TNFα signalling (p-value: 0.318; NES: −1.19) and TGF-β Signalling (p-value: 0.235; NES: −1.16), consistent with the inflammatory and fibrotic features exhibited following cryoinjury (Figure 7G). We also identified the unfolded protein response pathway (p-value: 0.074; Normalised enrichment score (NES): −1.40), a well-established component of the integrated stress response, as amongst the most enriched signatures following cryoinjury (Figure 7G and Supplementary Figure 8H). We compared the expression levels of atf4, an effector transcription factor from the unfolded protein response, during development and adulthood (Supplementary Figure 9A). We could observe a significant upregulation of atf4 and atf3 specifically upon injury, 1 dpci, compared to homeostatic conditions during development (7 dpf) or sham-operated animals (Supplementary Figure 9B). atf4 mRNA HCR staining revealed low atf4 expression levels in sham controls (Supplementary Figure 9C) and increase of its expression at the IA and injured border area at 1 dpci (Supplementary Figure 9D). Atf4 downstream targets involved in aminoacid synthesis, transport, and RNA charging were significantly upregulated at 1 dpci (Supplementary Figure 9E)^36^. These results show a plausible novel molecular target to modulate liver regeneration upon injury.

Many of the downregulated hallmarks and Gene Ontology (GO) enrichment (Figure 7G and Supplementary Figure 8G), which we observed following cryoinjury at 1dpci, are associated with housekeeping functions of hepatocytes, such as bile acid metabolism (p-value: 0.02; NES:1.87), xenobiotic metabolism (p-value: 0.037; NES:1.75), and fatty acid metabolism (p-value: 0.05; NES:1.65) (Figure 7G). At 3 dpci, 167 upregulated DEGs were detected, while 163 DEGs were downregulated (Figure 7C and E-F) compared to the sham. GSEA of DEGs at 3 dpci revealed several hallmarks related to cell proliferation and migration, including the G2/M checkpoint signature (p-value: 0.000; NES: −1.55) and epithelial-mesenchymal transition signature (EMT; p-value: 0.000; NES: −1.76) respectively (Figure 7G). These signatures are consistent with the timing of proliferation described following cryoinjury (Figure 5D), and the increased inflammation and recruitment of leukocytes to the injured area in livers between 3 dpci and 7 dpci (Figure 4D-F and Supplementary Figure 4D-F). To infer changes in tissue composition following cryoinjury, we used cell-type specific genes to identify the dynamics of liver regeneration on the different cell types present in the liver during the first 7 dpci (Figure 7B-D and 7H). At 1 and 3 dpci, we observed an increase in genes specifically expressed in ECs (Figure 7H), which may be indicative of a quick angiogenesis response as previously observed by confocal microscopy (Figure 6I-J), quickly reducing by 7 dpci when the ECs population seems almost fully recovered (Figure 6L and 6N). At 1 dpci, we detected an enrichment in genes expressed in hepatoblast, which may explain the dedifferentiation of hepatocytes to a hepatoblast-like state (Figure 7H). Genes associated with immune cells were enriched at 1 dpci upon injury, consistent with the early recruitment of immune cells towards the site of injury (Figure 7H). At 3 dpci, we detected enrichment in genes specifically expressed by BECs, suggesting an active restoration of the biliary network (Figure 7H). We detected an increase of HCs transcripts, showing the initial recovery of the two main liver parenchyma. HSCs were enriched at 3 dpci. This may explain the rapid development of the fibrotic response upon cryoinjury (Figure 7H). By 7 dpci, we detected 97 upregulated and 290 downregulated DEG compared to sham-operated livers (Figure 7D and E-F), suggesting a gradual shift towards homeostasis in the liver. When we compared the expression between the injured area in the ventral lobe with the contralateral lobe, we observed a much bigger proportion of DEG in the injured area when compared to the contralateral lobes (Supplementary Figure 8C-F). This might explain previous microscopy observations of a strong localised regeneration combined with a compensatory response in the livers during regeneration. In light of these studies, we conclude that the transcriptional landscape dynamically changes during regeneration as the liver responds to cryoinjury, undergoes regenerative growth and eventually returns to homeostasis.

## Discussion

Here, we introduce the hepatic cryoinjury model, which serves as a rapid and reliable method to investigate liver regeneration in a spatially localised manner (Figure 8 and Table 1). The frozen cryoprobe’s contact with the liver surface generates a necrotic region, surrounded by apoptosis in the injured border area, resembling clinical liver injuries^36^. Furthermore, necrotic cells trigger a robust innate immunological response and immune infiltration at the injury site. The hepatic cryoinjury model offers several advantages compared to the other injury models (Table 1). Firstly, it creates a distinct localized injury, enabling the examination of injured and uninjured hepatic parenchyma in the same animal. Moreover, hepatic cryoinjury is easy to perform and visualise macroscopically (blister formation), which facilitates the investigation of the tissue regenerative microenvironment and the utilization of state-of-the-art sequencing and imaging techniques. Secondly, the cryoinjury model gives rise to necrotic and apoptotic tissue, and the localised infiltration of inflammatory cells, which are absent in other surgical models (i.e., HPx). Thirdly, the hepatic cryoinjury model induces a transient fibrotic response not seen in other models, which resolves within 30 dpci without hindering liver repair. For these reasons, we anticipate that this unique model will aid in the discovery of the molecular mechanisms of fibrosis and regeneration.

**Table 1.** Comparison of liver injury and regeneration methods in animal models.

|  | Cryoinjury | HPx | Genetic ablation | APAP |
| --- | --- | --- | --- | --- |
| Tissue affected | Parenchymal and non-parenchymal cells | Apoptosis limited to the tissue around the amputation site | Promoter-specific ablation (Only hepatocytes) | Pericentral hepatocytes and surrounding parenchymal and non-parenchymal cells |
| Regeneration | 30 | 5-7 | 5 | 5-7 |
| Fibrosis | Yes | No | No | No |
| Localised injury response | Yes | No | No | No |
| Regeneration response | Local hyperplasia & Compensatory growth | Compensatory growth | BECs Transdifferentiation | Compensatory hyperplasia |
| Inflammation | Major | Major | Major | Minor |
| Specific transgene | No | No | Yes | Yes |

**Figure 8.**
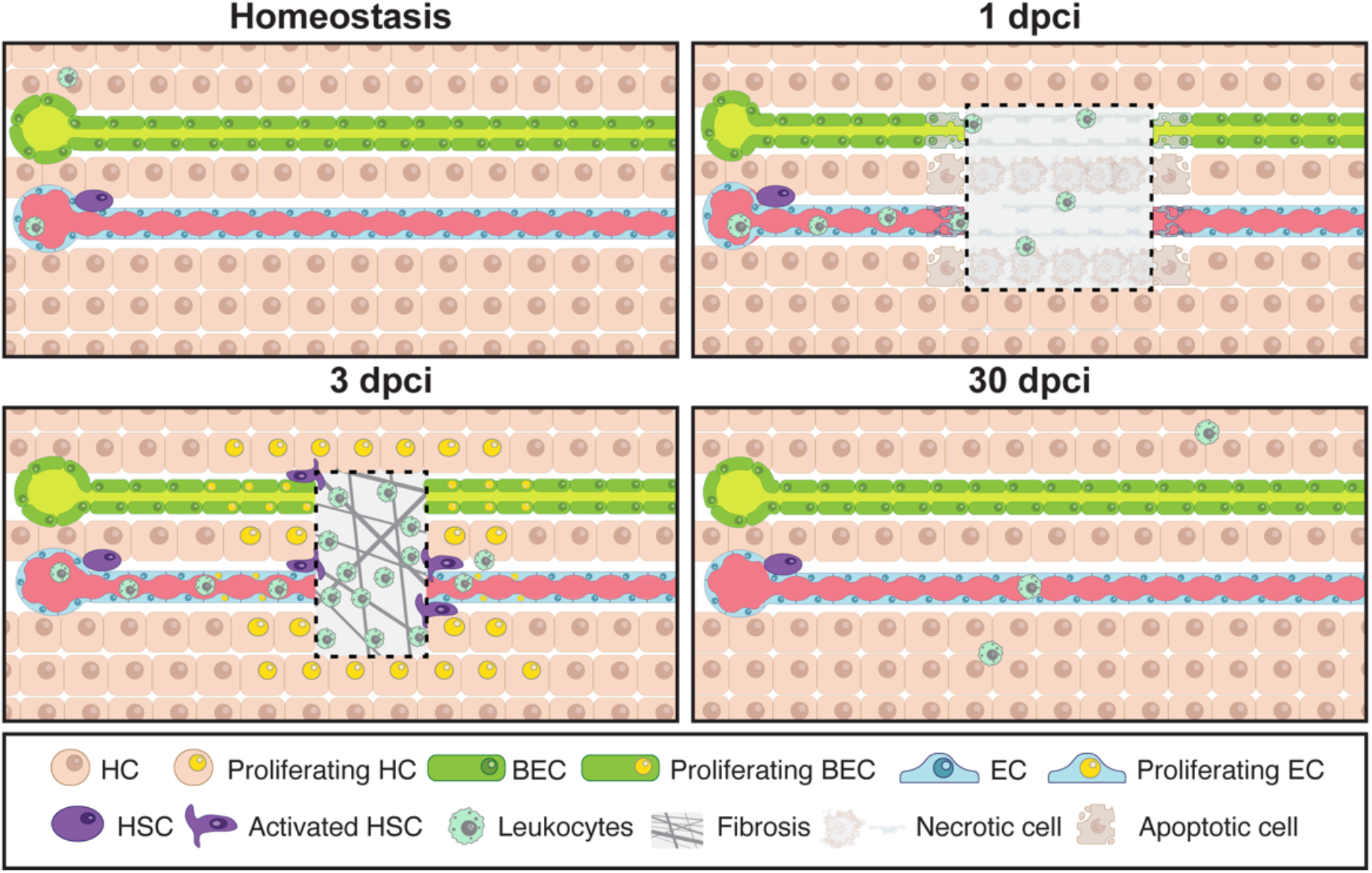
Model of liver regeneration upon cryoinjury. Schematic representation of the cellular events upon cryoinjury during liver repair in the zebrafish

Although the HPx model is almost 90 years old, it remains the gold standard of models to study liver regeneration. Liver regeneration in rodent HPx models occurs rapidly via compensatory hyperplasia, in which hepatocyte proliferation peaks at 36 and 48 hours post resection, before receding by 3 days and liver homeostasis is re-established at 7 days^1,37^. Zebrafish HPx models show similar dynamics of hepatocyte proliferation with almost no detectable proliferative beyond 4 days^7,8^. A major point of difference between the HPx models and hepatic cryoinjury is the more prolonged time course of regeneration (30 days). We suspect this is partly due to transient fibrosis and the need to resolve damaged tissue before mounting a proliferative phase of regeneration.

Acetaminophen (APAP) overdose is the most common cause of acute liver failure (ALF), as it induces necrosis of pericentral hepatocytes, leading to a necrotic area in zone 3 of the liver parenchyma^21^. The APAP model shares features of the hepatic cryoinjury model, such as the induction of region-specific necrosis (Table 1). However, one major distinction is that due to its mechanism of action, APAP targets pericentral hepatocytes, whereas the hepatic cryoinjury affects all cells present at the site of injury. Tissue loss due to APAP exposure is recovered by hepatocyte proliferation within 4 days, peaking at 2 days post APAP exposure^29^, which is similar to the kinetics of hepatocyte proliferation observed following HPx but earlier than we have observed following hepatic cryoinjury. Interestingly, hepatocyte proliferation following APAP exposure occurs in areas where healthy hepatocytes are in contact with necrotic tissue, rather than via compensatory hyperplasia^38^, which is reminiscent of the epimorphic proliferative response observed following cryoinjury. Another similarity between cryoinjury and APAP, is that APAP exposure triggers the infiltration of immune cells, activating an innate and adaptive immunological response^21^. Given the crucial role of immunity in liver disease, it is beneficial

Genetic ablation models provide an invaluable approach for targeted cell ablation in animal models, elucidating the roles of various cell populations in liver tissue repair and regeneration^39,40^ (Table 1). These models have significantly contributed to our understanding of transdifferentiation events between BECs and hepatocytes in the context of tissue regeneration. While the ability to selectively target Liver fibrosis is a critical feature in the development of chronic liver diseases and the progression towards hepatocellular carcinoma (HCC). Hepatic cryoinjury leads to a transient fibrotic response (peaking at 5 dpci) that was resolved within 30 dpci. Compared to other models of liver fibrosis (e.g., CCl_4_, TTA, ethanol or DDC), which typically develop over months^26^ (Table 1), fibrosis in the cryoinjury model occurs rapidly and is resolved over time. In future studies, the cryoinjury model could be used to explore the clonality of fibrosis and regeneration. To this end, recent advances in DNA-barcoding lineage tracing methods^41–45^ and spatial transcriptomics^46,47^ make it feasible to that the hepatic cryoinjury model exhibits classic hallmarks of inflammation specific cell populations presents clear advantages, generating cell type-specific promoter transgenic lines can be labour-intensive and time-consuming. In contrast to the cryoinjury model, which affects all parenchymal and non-parenchymal cells in the liver, cell-specific ablation techniques uniquely target distinct liver cell populations, thereby only allowing for the investigation of the function of a unique liver cell of interest. use the cryoinjury model to identify the mechanisms driving the resolution of fibrosis. This enhanced understanding is crucial for identifying innovative therapeutic strategies promoting liver repair and combating fibrotic progression.

In summary, we have described a new hepatic cryoinjury and regeneration model that recapitulates key features of liver disease (Figure 8). Given the rapid and reproducible nature of the technique, we suspect the model will offer opportunities to uncover the molecular underpinnings of liver regeneration.

## Material and methods

### Experimental Model and Details

Experiments were conducted with adult zebrafish (Danio rerio) aged 4-9 months raised at a maximal 5 fish/L and maintained under the same environmental conditions: 27.5-28C, 650-700ms/cm, pH 7.5. The lighting conditions were 14:10 hours (Light: Darkness) and 10% of water exchange a day. Experiments were approved by the Peter MacCallum Cancer Centre AEEC (E665). All animal procedures conformed in accordance with the Prevention of Cruelty to Animals Act, 1986 (the Act), associated Regulations (2019) and the Australian code for the care and use of animals for scientific purposes, 8th edition, 2013 (the Code). The following transgenic zebrafish lines were used in this study: *Tg(fabp10a:NLS-mCherry)*^53^, *Tg(fabp10a:GreenLantern-H2B)*^*uom306* 54^, *Tg(fabp10a:rtTA;TRE-Cre)*^*uom307*^ *herein referred to as Tg(fabp10a:Tet-ON-Cre), Tg(fabp10a:NLS-mKate), Tg(fli1a:nEGFP)*^*y7* 55^.

### Cryoinjury and Analysis of the Injured Area

Fish are immersed in Tricaine (0.032%; wt/vol; Sigma) before surgery and placed in a foam holder with the ventral area facing up. Under a stereo microscope, scales above the liver area are removed using sharp forceps. Once the area had been cleaned, a small incision was made parallel to the skin using micro-dissecting scissors towards the midline, and the anterior fin was made to expose the liver through the ventral zebrafish body. Once the liver is exposed, the excess water is removed using tissue paper (KimWipe). The cryoinjury probe of 1mm diameter (Jaycar; Australia), which is cooled in liquid nitrogen for one minute, is placed on the liver surface for 15 seconds until thawing. Sham operations consisted of placing the thawed cryoprobe on the exposed liver at room temperature. Fish are quickly placed in a freshwater tank and reanimated by pipetting water into their gills for 30 seconds. The complete surgical procedure takes 3-5 minutes per fish. Healthy fish should be swimming normally after 5 minutes. Fish are transferred to the normal tanks and put into their racks 10 minutes after surgery with a gentle water flow. The damaged tissue was easily identified either from the lack of fluorescence when using HCs specific promoter line, which expresses a nuclear mCherry or GreenLantern signal specifically in HCs, or a blister-like injury in the liver surface. Injured and SHAM livers were phenotyped, and acquisitions were performed using a Leica M165FC fluorescent stereomicroscope. The injured area was measured on Imaris Software 9.0 (Bitplane). Statistical analysis and graphs were generated using GraphPad Prism v9.

### Whole-Mount CUBIC Liver Immunofluorescence and Image Processing

*In toto* immunofluorescence was performed as described^48^. Animals were euthanized at different times post-injury by immersion in 0.16% Tricaine. Livers were collected and fixed in PFA 4% overnight at 4 degrees in rocking agitation. We washed the livers 3 times in PBS Tw 0.1% before transferring them to CUBIC-L^49^ for 18h at 37C. After CUBIC-L incubation, livers were washed in PBS1x three times for 15 minutes. Livers were incubated overnight with blocking solution (5% BSA, 5% goat serum, and 0.1% Tween-20). Primary and secondary antibodies were incubated for one day at 4C in rocking agitation. Livers were transferred to CUBIC-R^+^ for 2 hours at room temperature and embedded in CUBIC-R^+^-Agarose for imaging. Livers were mounted on a glass bottom culture dish (MatTek Corporation) for confocal acquisition. Whole liver acquisitions were obtained with Zeiss LSM 780 and Nikon Sora Spinning-Disk confocal microscopes using 10X dry and 20X dry lenses. Images were acquired at 512×512 or 1024×1024 resolution. For every liver acquisition, tile-scan and z-stack modules were used to acquire a representative area of the liver. The injured area of the liver in whole-mount acquisitions was analysed and measured on Imaris Software 9.0 (Bitplane) and Fiji^50^. Acquisitions were saved as TIFF or JPEG on Fiji of the original files.

### BrdU^+^ and PCNA^+^ hepatocytes Image Analysis

Adult zebrafish were injected intraperitoneally at 2 dpci with 20 μl of 2.5 mg/ml of 5-Bromo-2-deoxyuridine (BrdU, B5002-1G, Sigma). Livers were collected and processed for analysis at 5 dpci. *in toto* immunofluorescence was performed as described^48^. Whole-mount IF was performed using anti-BrdU (Abcam # ab6326 [BU1/75 (ICR1), rat), anti-RFP (Abcam #ab62341, rabbit), Goat anti-Rat IgG (H+L) Cross-Adsorbed Alexa Fluor 647 (ThermoScientific), and Goat anti-Rabbit IgG (H+L) Alexa Fluor 594 Antibody (Invitrogen). Using Imaris Software 9.0 (Bitplane), we used the co-localisation channel tool to create a new channel, which represents BrdU^+^ HCs. A spot segmentation of BrdU^+^ HCs and BrdU^-^HCs was created. Manually, the site of injury was created as a surface. We used the new injured area surface and the ImarisXT/Distance-Transformation package (outside surface) to define the distance of every BrdU^+^ HCs spot to the injury site. This new channel was masked into the BrdU^+^ HCs spots, which were previously created. The novel BrdU^+^ HCs spots channel contains the specific distance to the injured area from every proliferating HCs, named Proliferating HCs distance channel. A new spot segmentation was done using the Proliferating HCs distance channel. We exported the Position X, Y, Z and Proliferating HCs distance channel mean intensity from the last Proliferating HCs distance channel. Spot segmentation. Using R, the different datasets were selected to create a representative graph, using ggplot2 package, of the presence of BrdU^+^ HCs or PCNA^+^ HCs towards the site of injury and quantified their location throughout the parenchyma with the injured area as the point of reference, and the density towards the slide or whole-mount acquisitions.

### Clodronate intraperitoneal injections

Intraperitoneal (IP) injections of 10 μl PBS and clodronate liposomes (5 mg/ml) (C-005, Liposoma BV, Amsterdam, The Netherlands) were performed in each fish, one day before the cryoinjury in the adult zebrafish livers. Livers were collected with the rest of the gastrointestinal tract (GIT) and processed for analysis at 3 and 7 dpci. PBS and clodronate injected livers were phenotyped and acquired using a Leica M165FC fluorescent stereomicroscope. The injured area was measured using Fiji Image Analysis Software^50^. Statistical analysis and graphs were generated using GraphPad Prism v9. Livers were fixed overnight in 4% PFA for cryosectioning and immunostaining.

### Doxycycline treatment

Adult fish were treated by immersion in Doxycycline (30 mg/mL; Sigma D5207) for 72 hours. Animals were kept in darkness at 28C during the duration of the treatment. Animals were fed three times in a fish clean water and placed back in the Doxycycline-treated water.

### Zebrafish Liver Fixation and Histological Processing

To obtain histology sections for AFOG or immunofluorescence, adult zebrafish livers were collected with the rest of the GIT. This procedure minimized disruption of the fragile liver tissue. Adult zebrafish were euthanized by immersion in 0.16% tricaine and placed in ice-cold PBS. A ventral incision through the skin, muscle, and peritoneal cavity was made, starting at the anus towards the pericardiac cavity. To avoid disrupting adhesions and scar tissue at the injury zone in early injury time points (1-7 dpci), the tissue adhering to the liver injury was left intact on the liver by performing an incision surrounding it. The GIT was detached at its anal part and slowly lifted while cutting the vasculature connecting the dorsal part of the GIT to the peritoneum and gonads. Once completely lifted from the cavity, a final cut was made anteriorly on the oesophagus to extract the GI. Samples were fixed overnight at 4°C in 10% Neutral Buffered Formalin (HT501125, Sigma) and embedded in paraffin following standard processing (45 min. incubations in ethanol 70%, 90%, 95%, 2 x 100%, 2 x xylenes, and 4x paraffin washes). 7 µm sagittal sections were mounted on SuperFrost+ slides (#12-550-15, Fisherbrand) and dried overnight at 37°C before storage or processing.

### Acid Fuchsin-Orange G (AFOG) Staining and Imaging

For AFOG staining, sections were dewaxed and rehydrated using standard procedures and then stained as described^51^. Collagen was stained blue, while fibrin and cell debris appeared red. Sections were imaged on a Leica DM6-B upright microscope equipped with a Leica DFC7000 camera using the 10x or 20x objectives. Whole GIT images were acquired using the LAS-X software’s automated navigator and stitching function.

### Hematoxylin and Eosin (H&E) Staining and Imaging

For H&E staining, sections were sectioned at the 5 μm thickness. Slides were dewaxed and rehydrated using standard procedures. Slides were stained with Mayer’s Hematoxylin solution (ab220365; Abcam) for 5 minutes. The nuclear staining was differentiated for 15 seconds in 0.37% HCl prepared in 70% ethanol, and the slides were washed in tap water for 10 minutes. Sections were incubated on 0.1 % Eosin Y solution (ab246824; Abcam) for 3 minutes in 0.1% water with acidic acid. Sections were washed in double distilled water. The sections were dehydrated in a water/ethanol series, cleared in xylol, and mounted in DPX mounting medium (06522; Sigma-Aldrich). Sections were imaged on an Olympus SLIDEVIEW VS200 automated slide scanner using 10x, 20x, 40x and 63x lenses. Whole GIT sections were acquired, and images were analysed using QuPath^52^ and Olympus LS software.

### Immunofluorescence in Cryosections

Cryosections were incubated for 30 min in PBS at 37°C. Sections were washed three times on PBST (0.1% Tween 20 in PBS). Slides were permeabilised in 0.5% Triton X-100 in PBS for 15 minutes. Slides were washed three times for 5 min in PBST after permeabilization. After washing the slides in PBST and drawing hydrophobic rings (ImmEdge pen, H-4000, Vector), unspecific binding sites were blocked using 5% goat serum, 5% bovine serum albumin in PBST for 2 h at room temperature. Primary antibodies were incubated in blocking buffer overnight at 4°C in a humidity box. Primary antibodies used were, rabbit anti-mfap4 (GTX132692, GeneTex; 1:300), rabbit anti-mpx (GTX128379 GeneTex; 1:300), rabbit mCherry (ab167453, Abcam; 1:500), mouse anti-zebrafish gut secretory cell epitopes (ab71286, Abcam; 1:400), chicken anti-GFP (ab13970, Abcam; 1:300). Slides were washed three times in PBST before incubating with secondary antibodies and DAPI as nuclear counterstaining at room temperature for 45 minutes. Secondary AlexaFluor conjugated antibodies (ThermoScientific, 1:400) were used to detect isotype-specific primary antibodies and mounted with Vectashield reagent (Vector Laboratories).

### Immunofluorescence in Paraffin Sections

Immunofluorescence was performed as described^3^ with the following modifications. After dewaxing and rehydration, sections were heated and then pressure cooked for 4 min. in citrate unmasking solution (H3300, Vector, 100x dilution). Once cooled, sections were permeabilized for 10 min in methanol, washed in PBST (0.1% Tween 20 in PBS), and photobleached for 45 min to remove autofluorescence (PMID: 29993362). Slides were washed three times for 5 min in PBST before incubation in 0.25% Triton X-100 in PBS. After washing the slides in PBST and drawing hydrophobic rings (ImmEdge pen, H-4000, Vector) unspecific binding sites were blocked using 5% goat serum, 5% bovine serum albumin in PBST for 1 h at room temperature. Primary antibodies were incubated in blocking buffer overnight at 4°C in a humidity box. Primary antibodies used were, mouse anti-RFP (clone RF5R, MA5-15257 ThermoFisher; 1:1000), mouse anti-PCNA (SC-56, Santa Cruz Biotechnology; 1:500), Rabbit anti-Lcp1 (GTX124420, GeneTex; 1:500), mouse anti-col1a1 (clone Sp1.d8, DSHB; 1:20), and mouse anti-Anxa4 (2F11, ab71286, Abcam). Slides were washed three times for 5 min in PBST before incubation with secondary antibodies and DAPI as nuclear counterstaining in blocking buffer for 1 h at room temperature. Alexa conjugated secondary antibodies (Life Technologies, 1:500) were used to detect isotype-specific primary antibodies, except for the detection of Col1a1, which required a tyramide amplification step (Alexa Fluor 555 Tyramide SuperBoost Kit, Invitrogen). After staining, sections were quickly rinsed in double distilled water and mounted with Fluorsave reagent (345789, Millipore).

### AFOG Histological Imaging and Analysis

After immunofluorescence, sections were imaged on a Zeiss LSM900 confocal microscope using the 10x or 20x objectives with 3-4 images per z-stack. Samples were imaged using the ZEN-pro software, and images were saved as multi-stack CZI files. Images were stacked in ImageJ, and individual channels were exported to TIF files. Multi-colour images and color levels were adjusted using Adobe Photoshop, and figure panels were cropped and rotated to dimension in CorelDRAW.

To quantify hepatocyte proliferation, sections from *fabp10a:n-m*Kate livers were immunostained with anti-RFP and anti-PCNA antibodies (see above). Three sections containing the largest injury areas were imaged. n-mKate^+^ and PCNA^+^n-mKate^+^ cells were counted manually using ImageJ in defined regions that include 150 µm adjacent to the border zone. The percentages of PCNA^+^n-mKate^+^/n-mKate^+^ cells from individual sections were averaged to establish the hepatocyte proliferation index for each animal.

To quantify the proliferation of the biliary epithelial cells, adjacent sections were immunostained with anti-Anxa4 and anti-PCNA antibodies (see above). Three sections containing the largest injury areas were imaged. Anxa4^+^ and PCNA^+^Anxa4^+^ cells were counted manually using ImageJ in defined regions that include 150 µm adjacent to the border zone. The percentages of PCNA^+^Anxa4^+^/ Anxa4^+^ cells from individual sections were averaged to establish the hepatocyte proliferation index for each animal.

To quantify leukocyte infiltration, sections from *fabp10a:n-mKate* livers were immunostained with anti-RFP and anti-Lcp1 antibodies (see above). Three sections containing the largest injury areas were imaged. The number of Lcp1^+^ cells was counted manually using ImageJ in defined regions (150 µm x 150 µm) that include the injury area and border zone. Lcp1+ counts in three independent regions per animal were averaged to establish a quantification for each animal.

To quantify collagen deposition, liver sections were immunostained using an anti-Col1a1 antibody and imaged using confocal microscopy. Three sections containing the largest injury areas were imaged. The Col1a1^+^ area/total liver parenchyma area ratio in three independent regions per animal was averaged to establish a quantification for each animal.

### Quantification and Statistical Analysis

Sample sizes were chosen based on previous publications and are indicated in each figure legend. No animal or sample was excluded from the analysis unless the animal died during the procedure. The experiments were not randomized, and the investigators were not blinded to allocation during experiments and outcome assessment. All statistical values are displayed as mean ± standard deviation. The figures or figure legends indicate sample sizes, statistical tests, and P values. Data distribution was determined before using parametric or non-parametric statistical tests. Statistical significance was assigned at P < 0.05. All statistical tests were performed using Prism 7 software.

### HCR RNA-FISH staining

Samples were fixed for 24h in PFA 4% at 4°C. Livers were washed 3 times for minutes in PBST (0.1% Tween 20 in PBS) on ice and permeabilised in 0.5% Triton X-100 in PBS rocking overnight. Livers were washed 2 times for 5 minutes on PBST and postfixed with 4% PFA for 20 minutes at room temperature. Livers were washed 3 times for 5 minutes on PBST. Samples were incubated in 1 mL probe hybridization buffer for 5 minutes. Afterwards, livers were incubated in 1 mL 37°C probe hybridization buffer for 30 minutes. Livers were incubated in 4 pmol probes overnight at 37°C rocking. Livers were washed 4 times by 15 minutes with wash buffer at 37°C and washed again in 5X SSCR twice for 5 minutes. Samples were pre-amplified with 250 µl of amplification buffer for 10 minutes at room temperature. Samples were incubated with 6 µl of hairpin by 100 µl of amplification buffer. Excess hairpins were removed by washing twice for 5 minutes with 5x SSCT. Samples were nuclei counterstain with DAPI in 5X SSCT (1:1000) for 45 minutes, and final wash of 15 minutes in 5x SSCT. Samples were mounted for imaging on a glass bottom culture dish (MatTek Corporation) or storage at 4°C protected from light before imaging. Two probes were designed, atf4a and fabp10a for HCR RNA-Fish. Samples were scanned using Nikon SoRa Spinning Disk Confocal with a 10X and 20X dry lenses. Acquisitions were saved as TIFF or JPEG on Fiji of the original files.

### Bulk RNA-seq

Sham and injured livers at 1, 3, and 7 dpci were phenotyped under the fluorescent stereomicroscope (NSZ-606 Binocular Zoom fitted with a NightSea SFA light base) to confirm the presence of insult upon cryoinjury. In addition, 3 adult zebrafish livers were pooled per tube, discriminating between the injured border and liver tissue from other lobes. Finally, 3 replicates of 3 pooled livers were used for library preparation. Livers were transferred to a final volume of 300uL of cold TRIzol™ (Thermo Fisher Scientific) per tube on ice. Livers were homogenized using the mechanical homogenizer for 30s on ice, with a plastic pestle, to ensure fine homogenization. RNA was extracted according to the manufacturer guidelines (Direct-zolT^M^ RNA MiniPrep kit, Zymo Research). RNA quality was confirmed using an Agilent 4200 Tapestation System. Libraries were sequenced in Illumina NextSeq 500, with paired-end 75bp reads to a depth of 15M reads per sample. 7 dpf larvae Bulk RNA-seq embryo liver datasets were obtained from public GEO accession number GSE19571156^61^.

### Bioinformatic Analysis

BCL files were converted to FastQ files on Galaxy using bcl2fastq2 (v2.20.0.422 – Illumina). Reads were mapped to the reference genome (Ensembl build 11, release 94) using Hisat2, version 2.1.0 (Kim et al., 2015), and counting was performed using featureCounts, version1.6.0 (Liao et al., 2014. For bioinformatics analysis, we compared SHAM, 1, 3 and 7 dpci livers. Downstream analysis was performed in R, version 4.2.2 (R Core Team, 2018). Counts were normalized and differential expression between design groups was tested using the package EdgeR. Principal component analysis (PCA) plots, volcano plots and heatmaps were generated using the ggplot2 package, version 3.4.0 (H. Wickham. ggplot2: Elegant Graphics for DataAnalysis. Springer-Verlag New York, 2016.). For the enrichment, we selected all the Gene Stable IDs, translated them to Mus musculus Gene Stable IDs, and obtained the ENTREZIDs and SYMBOLs using biomaRt package (Durinck et al., 2005). Statistically significant genes were defined with the following thresholds, log2FC≥1.5 and p-value ≤0.05. Transcriptional downstream signatures were examined using Gene Set Enrichment Analysis (GSEA v4.0.3) (Subramanian et al., 2005). Gene ontology analysis was done using Gene Ontology Panther (http://geneontology.org). Data is available under the GEO accession number GSE245878.

## Abbreviations

AFOG: Fuchsin Orange G
ALF: Acute liver failure
APAP: Acetaminophen (N-acetyl-para-aminophenol
BECs: Biliary epithelial cells
BrdU: 5-Bromo-2 -deoxyuridine
CCl4: Carbon tetrachloride
DAPI: 4′,6-diamidino-2-phenylindole
DDC: 3,5-diethoxycarbonyl-1,4-dihydrocollidine
DEG: Differentially expressed genes
DMN: Dimethyl nitrosamine
Dpci: Days post-cryoinjury
ECM: Extracellular matrix
EC: Endothelial cell
FFPE: Formalin-fixed, paraffin-embedded tissue
GIT: Gastrointestinal tract
GSEA: Gene Set Enrichment analysis
HCC: Hepatocellular carcinoma
HPx: Hepatectomy
HSCs: Hepatic Stellate cells
IA: Injured area
IP: Intraperitoneal
Mpci: Months post-cryoinjury
NES: Normalised enrichment score
NTR: Nitroreductase
PCNA: Proliferating cell nuclear antigen
PFA: Paraformaldehyde
TAA: Thioacetamide
Ypci: Years post-cryoinjury

## Acknowledgements

M.S.-M. is supported by the Early.Postdoc Mobility fellowship from the Swiss National Science Foundation (SNSF).

D.B received support from the Wallonie Bruselles International Postdoctoral Fellowship.

J.M.G.-R. is supported by the National Institutes of Health (1R01HL164749-01), the American Heart Association (19CDA34660207), the Corrigan Minehan Foundation (SPARK Award), and the Hassenfeld Foundation (Hassenfeld Research Scholar Award).

A.G.C is supported by an NHMRC Investigator Grant (GNT1176650), and an ARC Discovery Project Grant (DP200102693).

We also extend our thanks to the Peter MacCallum Cancer Centre Core Facilities and their staff who supported this work, namely the Centre for Advanced Histology & Microscopy, the Molecular Genomics Core, Flow Cytometry and Bioinformatics Core Facilities. We also thank the staff involved at the University of Melbourne Zebrafish Core Facility (DRUM) and Biology Optical Microscopy Platform (BOMP). We also thank the MCRI Bioinformatics Unit, which was supported by the Victorian Government’s Operational Infrastructure Support Program Finally, we thank members of the Cox Laboratory (Peter MacCallum Cancer Centre) González-Rosa Laboratory and the Organogenesis and Cancer Program at the Peter MacCallum Cancer Centre for the helpful discussions.

## Supplementary Figures

**Supplementary Figure 1.**
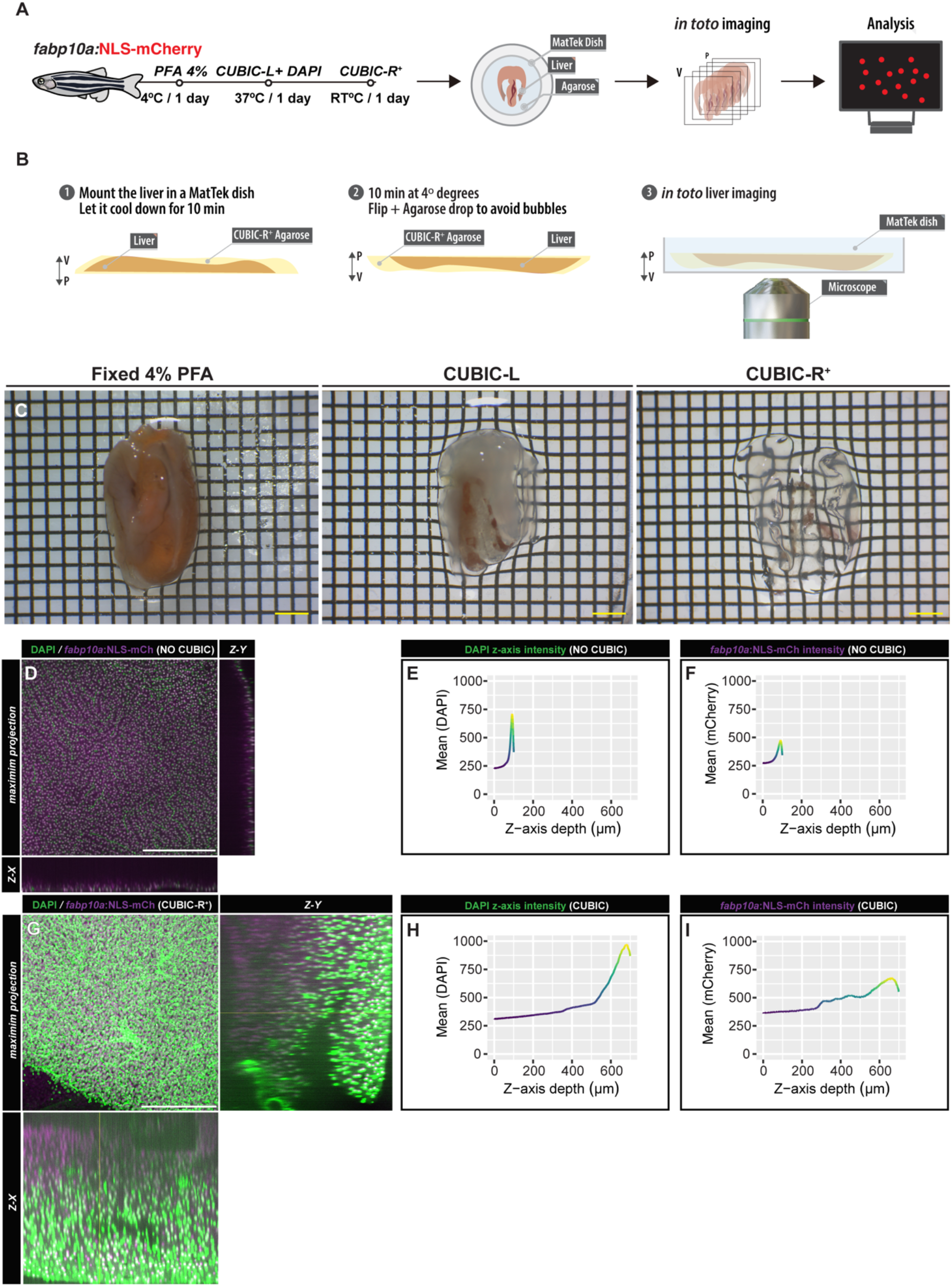
CUBIC-clearing liver method. A) Schematic representation of the liver CUBIC-clearing procedure for imaging the liver. B) Mounting strategy for imaging cleared livers. Livers are embedded in CUBIC-agarose within a MatTek dish, positioning the IA facing-up. The CUBIC-agarose disc containing embedded livers is cooled down for 10 minutes at 4 degrees Celsius. Once the CUBIC-agarose disc solidifies, it is carefully flipped using forceps, and a drop of CUBIC-agarose is added to prevent bubble formation. Cleared livers are prepared for imaging from this point onwards. c) An example of the liver-clearing process during the CUBIC-clearing procedure. D and G) z-stack representative examples, with Z-X and Z-Y projections, of non-cleared (D) and CUBIC-cleared (G) *in toto* liver confocal acquisitions. E-F) Mean intensity of DAPI (E) and mCherry (F) along the z-axis in non-cleared *in toto* liver confocal acquisition. H-I) Mean intensity of DAPI (H) and mCherry (I) along the z-axis in CUBIC-cleared *in toto* liver confocal acquisition. Scale bars: 2 mm (yellow), 500 µm (white).

**Supplementary Figure 2.**
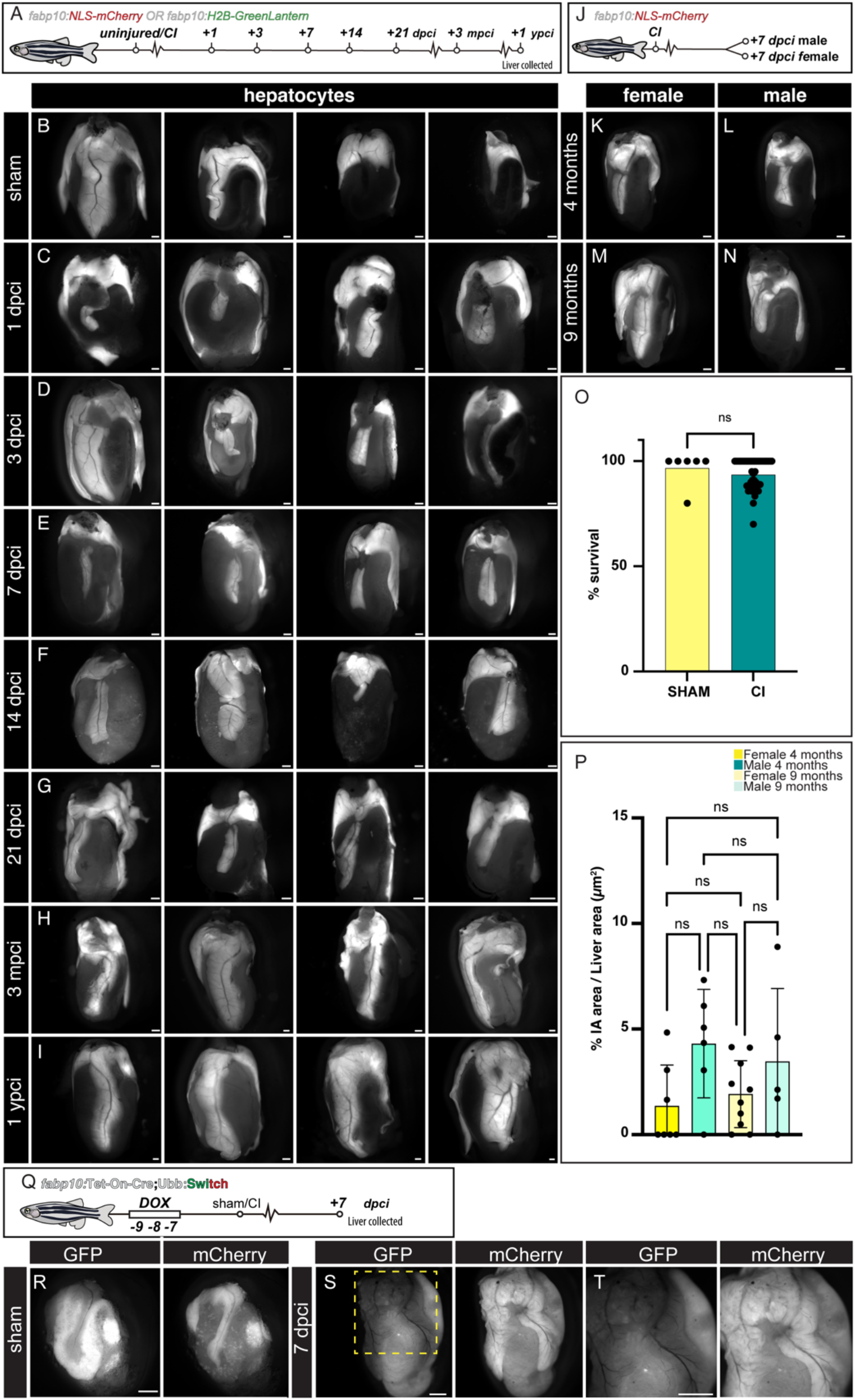
Temporal dynamics of liver regeneration in zebrafish following cryoinjury. A) Schematic representation of the timing of sample collection following cryoinjury. B-I) Epifluorescence acquisitions of adult zebrafish Tg(*fabp10a: GreenLantern-H2B*) or Tg(*fabp10a:NLS-mCherry*) livers at different stages of regeneration upon cryoinjury. J) Schematic representation of the timing of sample collection following cryoinjury between female and male adult zebrafish. K-N) Epifluorescence acquisitions of Tg(*fabp10a:NLS-mCherry*) livers in adult zebrafish differentiate between females (K and M) and males (L and N), as well as between those aged 4 (K-L) months and 9 months (M-N), at 7 dpci. O) Zebrafish survival upon cryoinjury (n= 444; survival= 92.97 %; *p-*value: 0.5843 unpaired Student’s *t*-test). P) Quantification of the IA area (n= 6, 4, 8, 5), bars indicate mean and SD, *p*-values: one-way ANOVA followed by Tukey’s multiple comparisons test). Q) Schematic representation of the timing of recombination and collection following cryoinjury. R-T) Epifluorescence acquisitions of adult zebrafish Tg(*fabp10a: Tet-ON-Cre; Ubb:Switch*) at sham and 7 dpci. CI: cryoinjury; Dox: Doxycline; dpci: day post-cryoinjury; mpci: months post-cryoinjury; ypci: years post-cryoinjury. Scale bars: 500 µm (B-I and K-N) and 1000 µm (R-T).

**Supplementary Figure 3.**
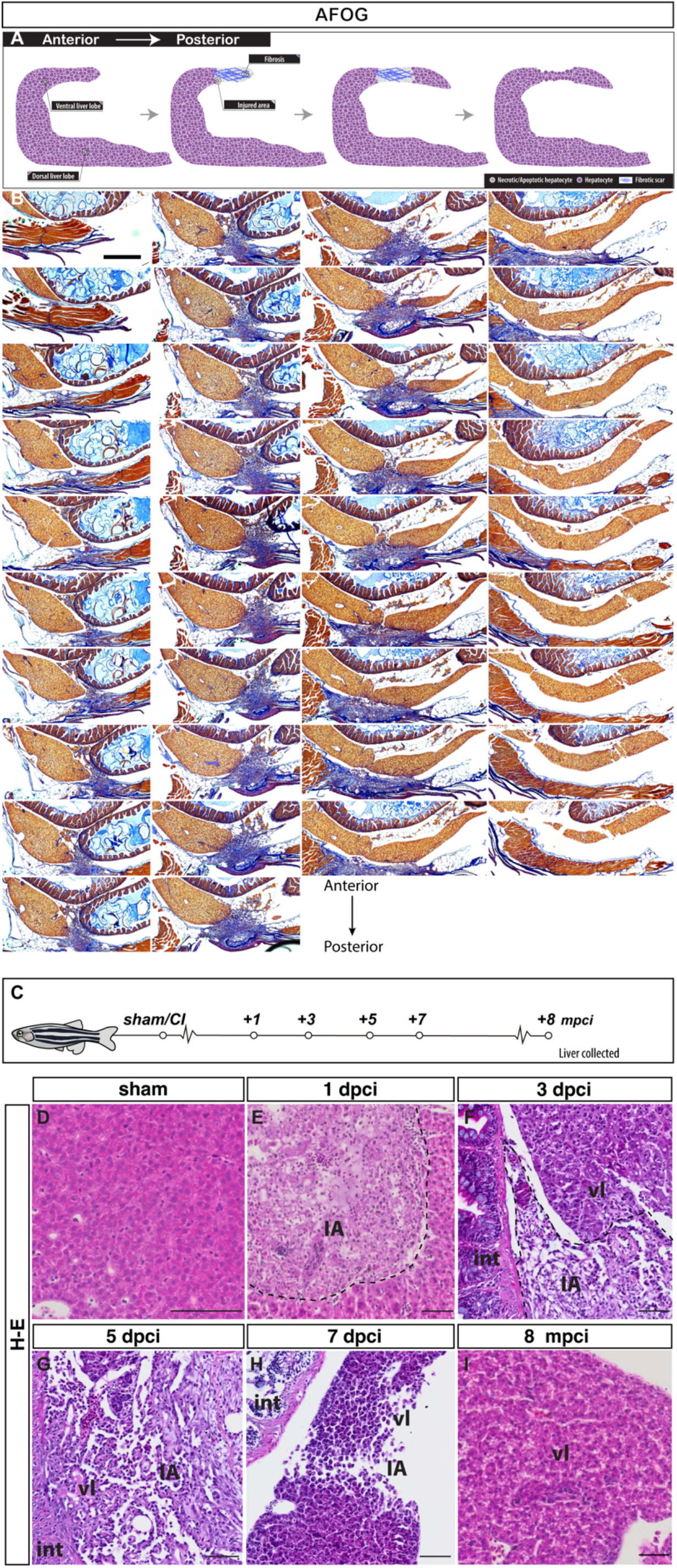
AFOG staining and histological appearance of regenerating livers following cryoinjury. A) Schematic representation of the histology section plane across 5 dpci adult zebrafish liver. B) AFOG staining of consecutive sections of 5 dpci adult zebrafish liver. Anterior is towards the left, dorsal is towards the top. C) Representation of the timing of sample collection following cryoinjury for histological examination. D-I) H&E staining of adult zebrafish liver, sham (D), 1 day post-cryoinjury (dpci; E), 3 dpci (F), 5 dpci (G), 7 dpci (H), and 8 months post-cryoinjury (mpci; I). Blue: collagen; Red: cell debris and fibrin; IA: injured area; int: intestine; vl: ventral lobe. Scale bars: 500 µm (B) and 50 µm (D-H).

**Supplementary Figure 4.**
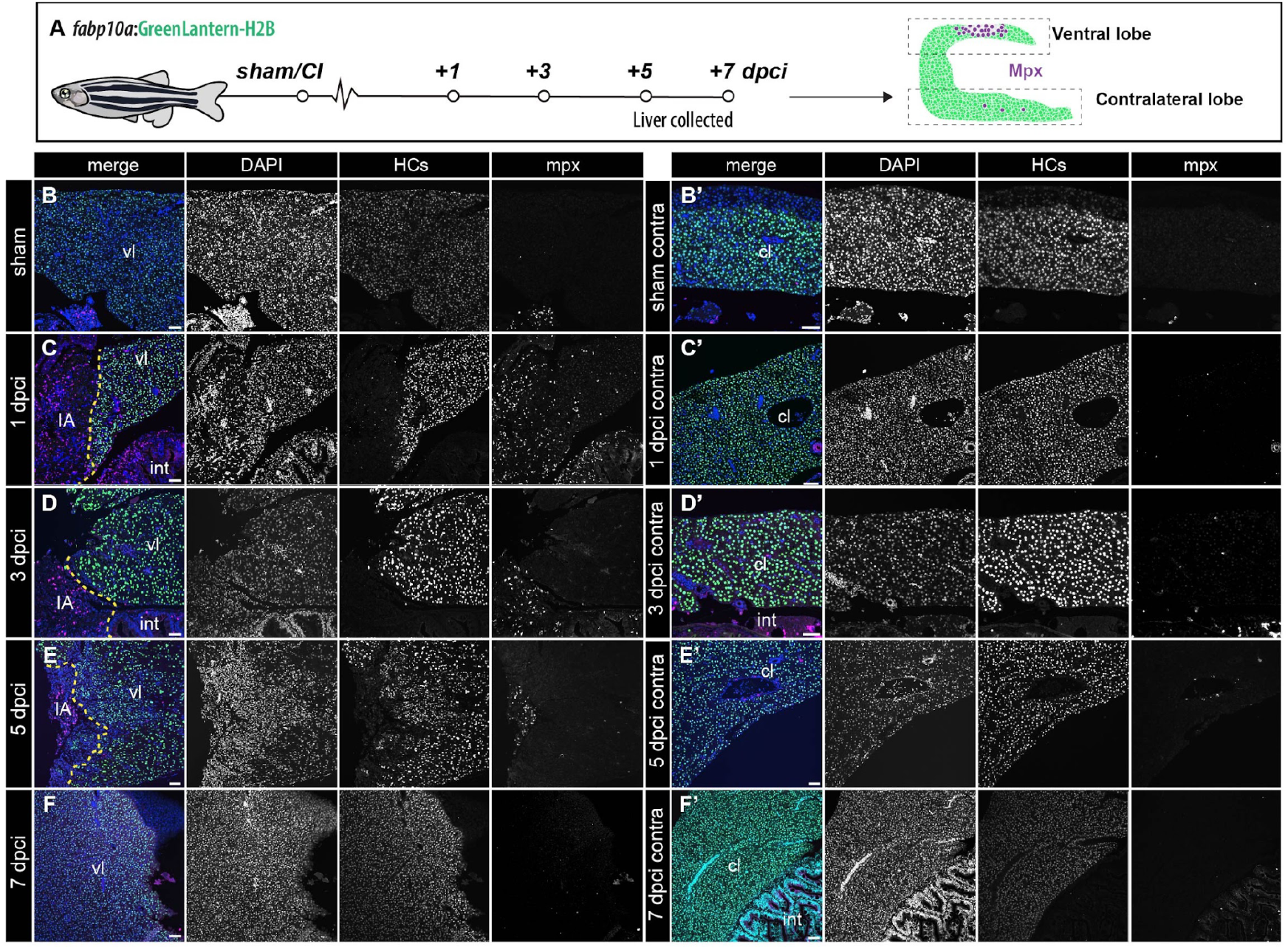
Mpx^+^ Neutrophils dynamics during liver regeneration upon cryoinjury. A)Schematic representation of the experiment workflow. B-F) Sections of livers from *Tg(fabp10a*:GreenLantern-H2B) animals at the indicated stages in ventral lobes compared to contralateral ones, immunostained to detect hepatocyte nuclei (GFP) and leukocytes (Mpx). cl: contralateral lobe; Dashed yellow line: border zone; IA: injured area. Scale bars: 50 µm.

**Supplementary Figure 5.**
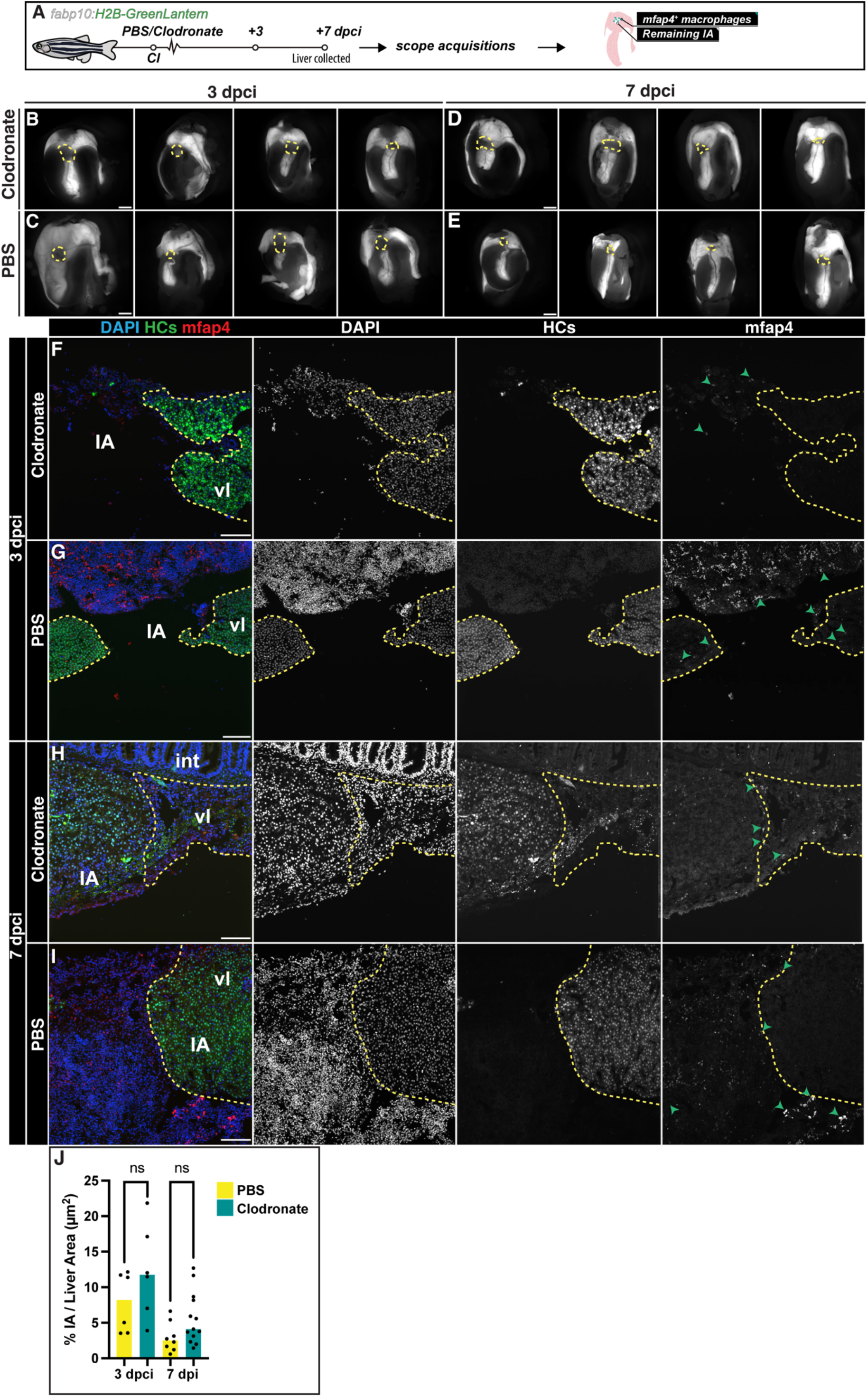
Clodronate effects on macrophages dynamics during liver regeneration upon cryoinjury. A) Schematic representation of the timing of clodronate injection and sample collection following cryoinjury. B-E) Epifluorescence acquisitions of adult zebrafish of Tg(*fabp10a:*H2B-GreenLantern) livers at 3 dpci with clodronate and controls with PBS IP injections (B-C), and 7 dpci with clodronate and PBS IP injections (D-E). F-I) Adult liver sections from cryoinjured animals at 3 dpci (F-G) and 7 dpci (H-I), immunostained to detect HCs nuclei (GreenLantern) and macrophages (mfap4) in clodronate injected samples (F and H), and PBS injected controls (G and I). J) Quantification of the IA area (n= 6, 6, 8, 13), bars indicate median, *p*-values: one-way ANOVA followed by Tukey’s multiple comparisons test. Dashed line: Border zone of the injured area; IA: injured area; int: intestine; vl: ventral lobe. Scale bars: 500 µm (B-E), and 100 µm (F-I).

**Supplementary Figure 6.**
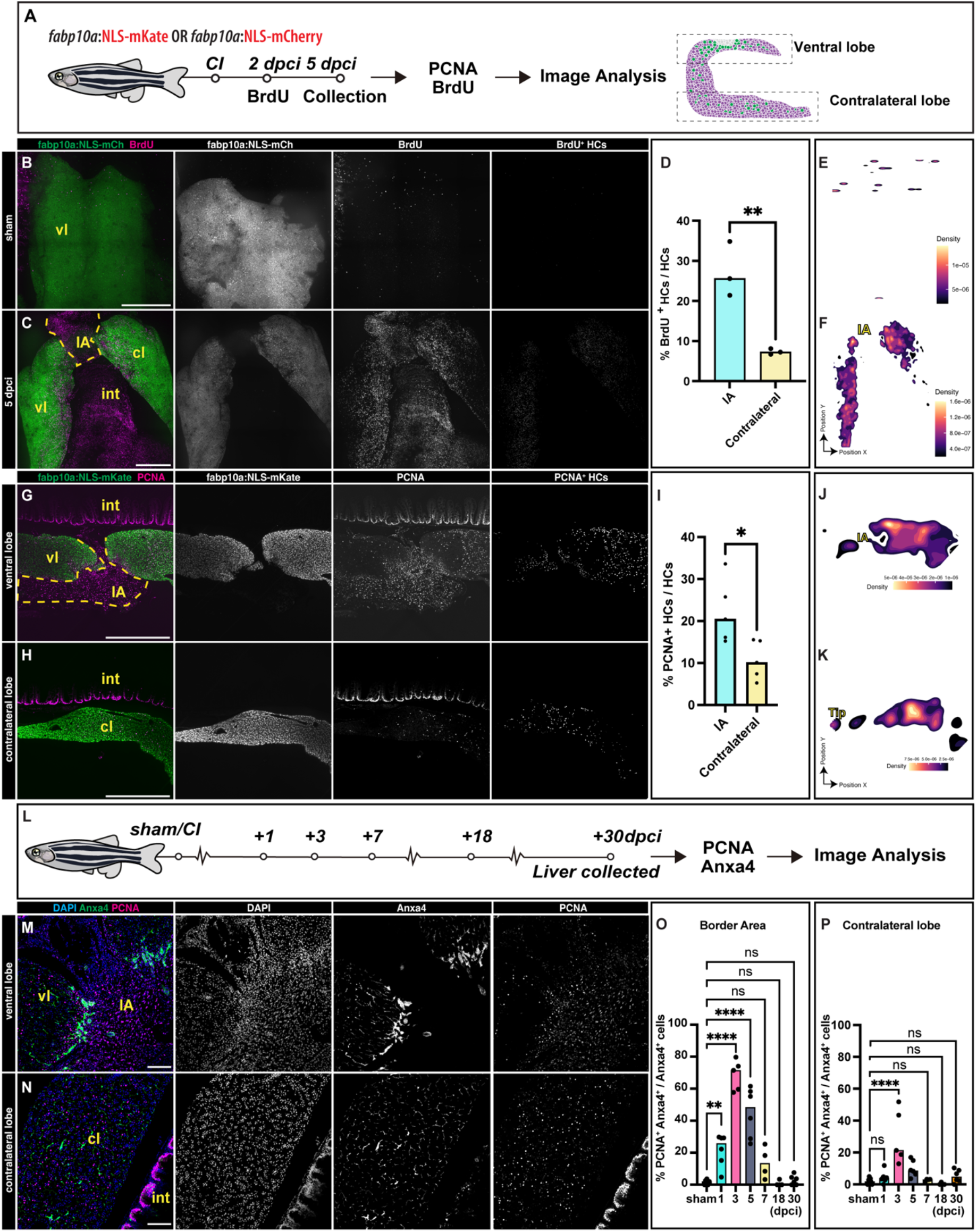
Cryoinjury induces local and distal compensatory hyperplasia. A) Schematic representation of the timing of sample collection following cryoinjury for BrdU and PCNA analysis. B) *in toto* of BrdU-pulse labelling sham adult zebrafish confocal acquisition. C) *in toto* of BrdU-pulse labelling 5 dpci adult zebrafish confocal acquisition. D) Quantification of BrdU^+^ mCherry^+^ HCs vs total number of mCherry^+^ HCs in the ventral lobe and contralateral one, bars indicate median, *p-*value: unpaired Student’s *t*-test. E-F) *in toto* confocal acquisition segmentation of BrdU^+^/mCherry^+^ HCs nuclei and density analysis using a Kernel Density Estimation (KDE) algorithm presented as a corresponding heatmap and quantification of the proliferating HCs density in sham animals (E) and 5 dpci (F). G) *in toto* of PCNA stained 5 dpci adult zebrafish confocal acquisition at the ventral lobe. H) *in toto* of PCNA stained 5 dpci adult zebrafish confocal acquisition at the contralateral lobe. I) Quantification of PCNA^+^ mKate^+^ HCs vs total number of mKate^+^ HCs in the ventral lobe and contralateral one, bars indicate median, *p-*value: unpaired Student’s *t*-test. J-K) *in toto* confocal acquisition segmentation of PCNA^+^/mKate^+^ HCs nuclei and density analysis using a Kernel Density Estimation (KDE) algorithm presented as a corresponding heatmap and quantification of the proliferating HCs density in the ventral lobe (J) and contralateral lobe (K). L) A simplified schematic illustrating the collection of livers following cryoinjury for PCNA^+^ and Anxa4^+^ biliary epithelial cells (BECs). M-N) Adult liver sections from cryoinjured animals at 5 dpci, immunostained to detect proliferation (PCNA) and BECs (Anxa4). O-P) Quantification of PCNA^+^ Anxa4^+^ BECs vs total number of BECs in the injured border area (O) vs contralateral lobe (P) from the indicated cohorts, (n = 6, 5, 5, 6, 4, 4 and 6; error bars representing SD; *p*-values: one-way ANOVA followed by Tukey’s multiple comparisons test). Dashed line: Border zone of the injured area; dpci: days post-cryoinjury; IA: injured area; int: intestine; vl: ventral lobe. Scale bars: 500 µm.

**Supplementary Figure 7.**
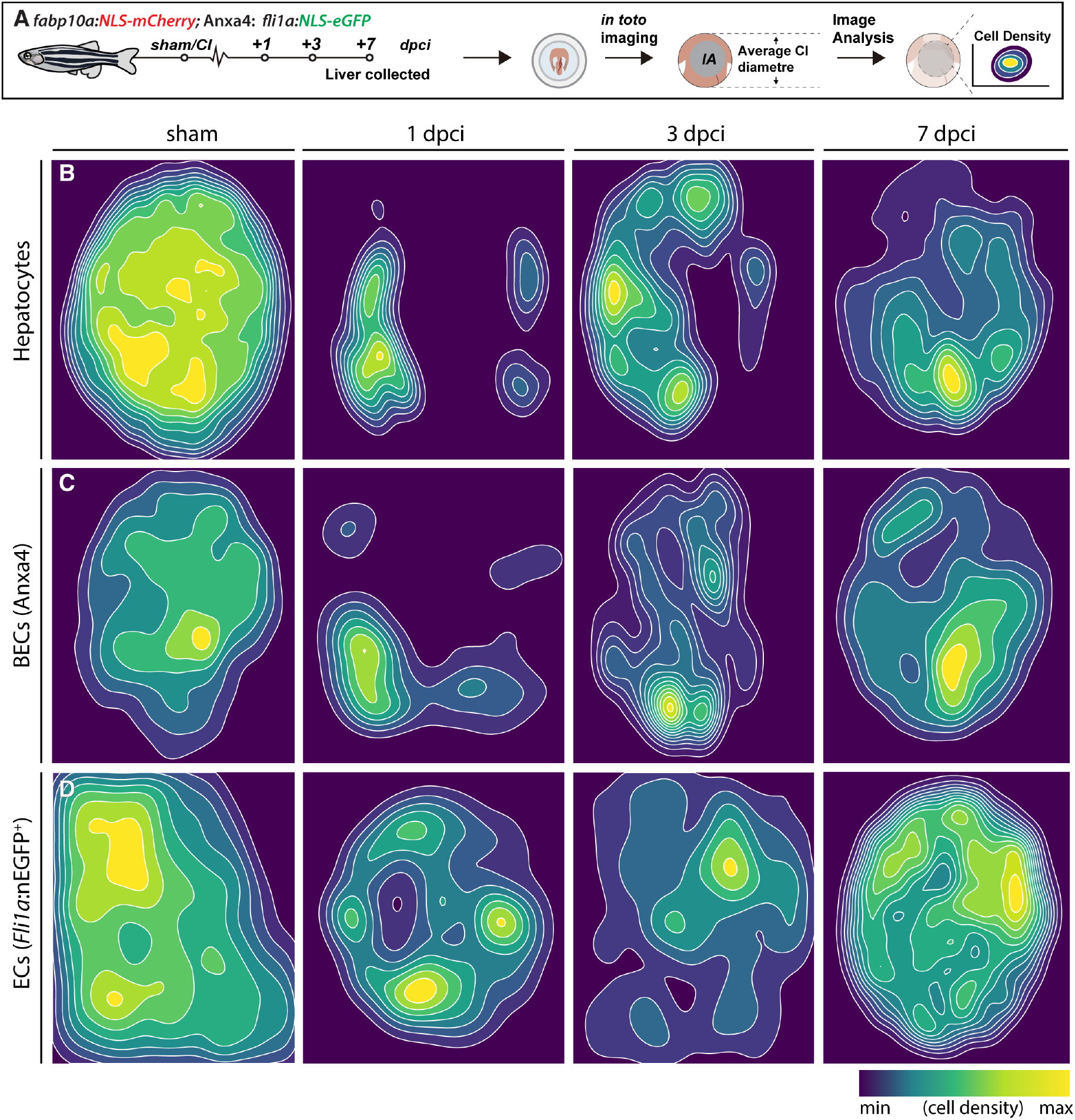
Regeneration of the injured area. **A**) Schematic workflow from data acquisition to image analysis. B-D) Region of interest (ROI) represents the integration of four independent datasets within the average injured area size of adult livers. (B) Tg(*fabp10a*:NLS-mCherry; n: 4), (C) immunostained against Anxa4 (n: 4), (D) Tg(*fli1a*:NLS-eGFP; n: 4) zebrafish. The probabilistic location of HCs (B), BECs (C), and ECs (D) is represented as a density plot over the first seven days of liver repair upon cryoinjury.

**Supplementary Figure 8.**
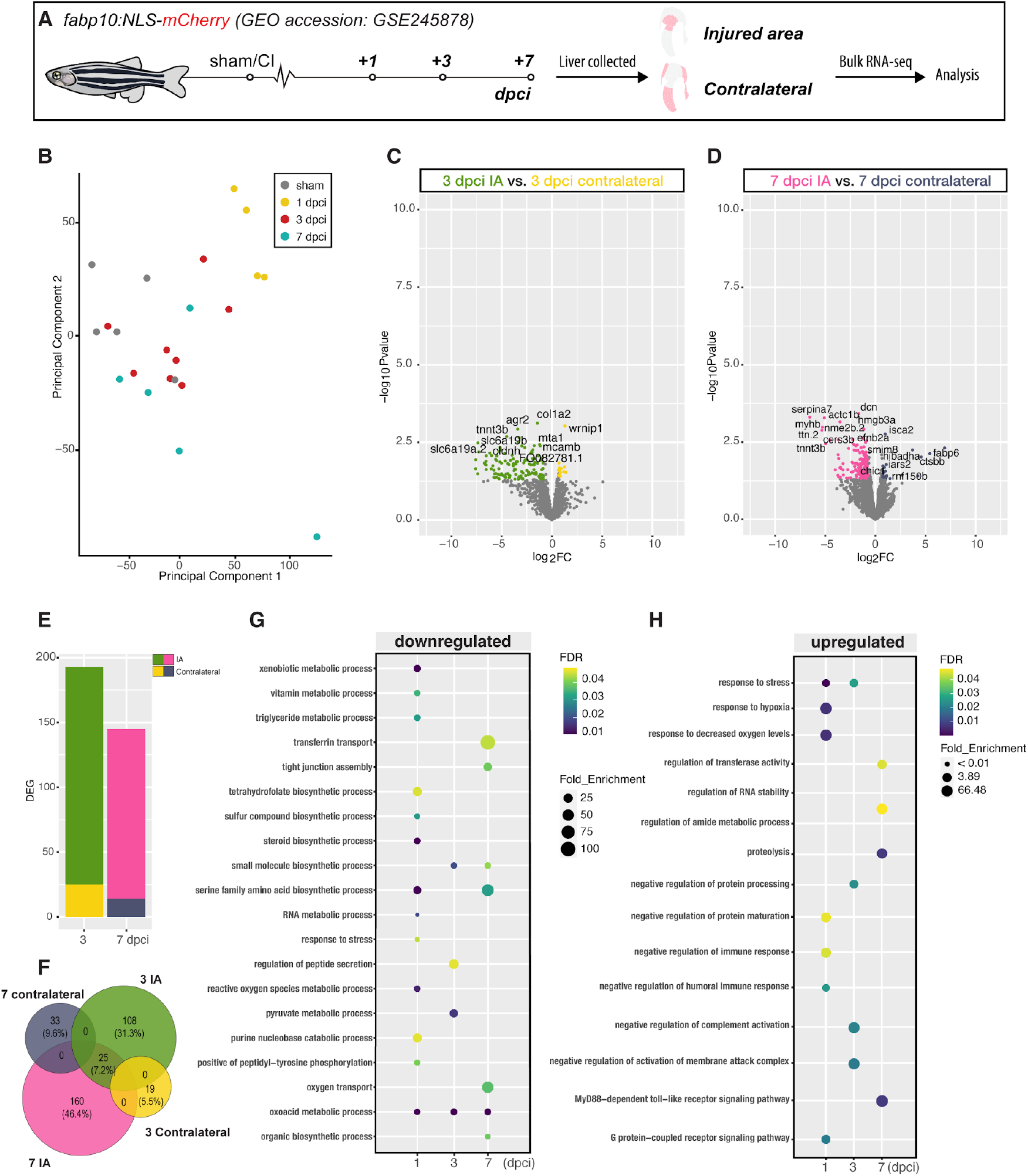
Comprehensive Gene Ontology (GO) Analysis of Regenerating Liver Tissue. A) Graphical illustration of the collection process (IA or Contralateral lobes) for liver repair stages following cryoinjury and sequencing. B) Principal Component Analysis (PCA) of sham-operated and injured ventral lobes at different stages including sham, 1, 3, and 7 dpci (Technical replicates = 4 per timepoint; livers = 16 per timepoint). C-D) Volcano plot representing Bulk RNA-seq data from the IA compared to Bulk RNA-seq data from contralateral liver lobes at 3 (C) and 7 (D) dpci. DEGs (FC ≥1.5 (Yellow or Navy Blue) or ≤-1.5 (Green or Pink); *P*≤0.05) with top DEG annotated. E) Bar plot representing the number of upregulated and downregulated DEG in IA and contralateral lobes at 3 and 7 dpci. F) Venn diagram representing DEG in IA and contralateral lobes at 3 and 7 dpci. G-H) GO-enrichment analysis of IA Bulk RNA-seq data showing both downregulated (G) and upregulated (H) terms during liver regeneration following cryoinjury in the IA of the ventral lobe.

**Supplementary Figure 9.**
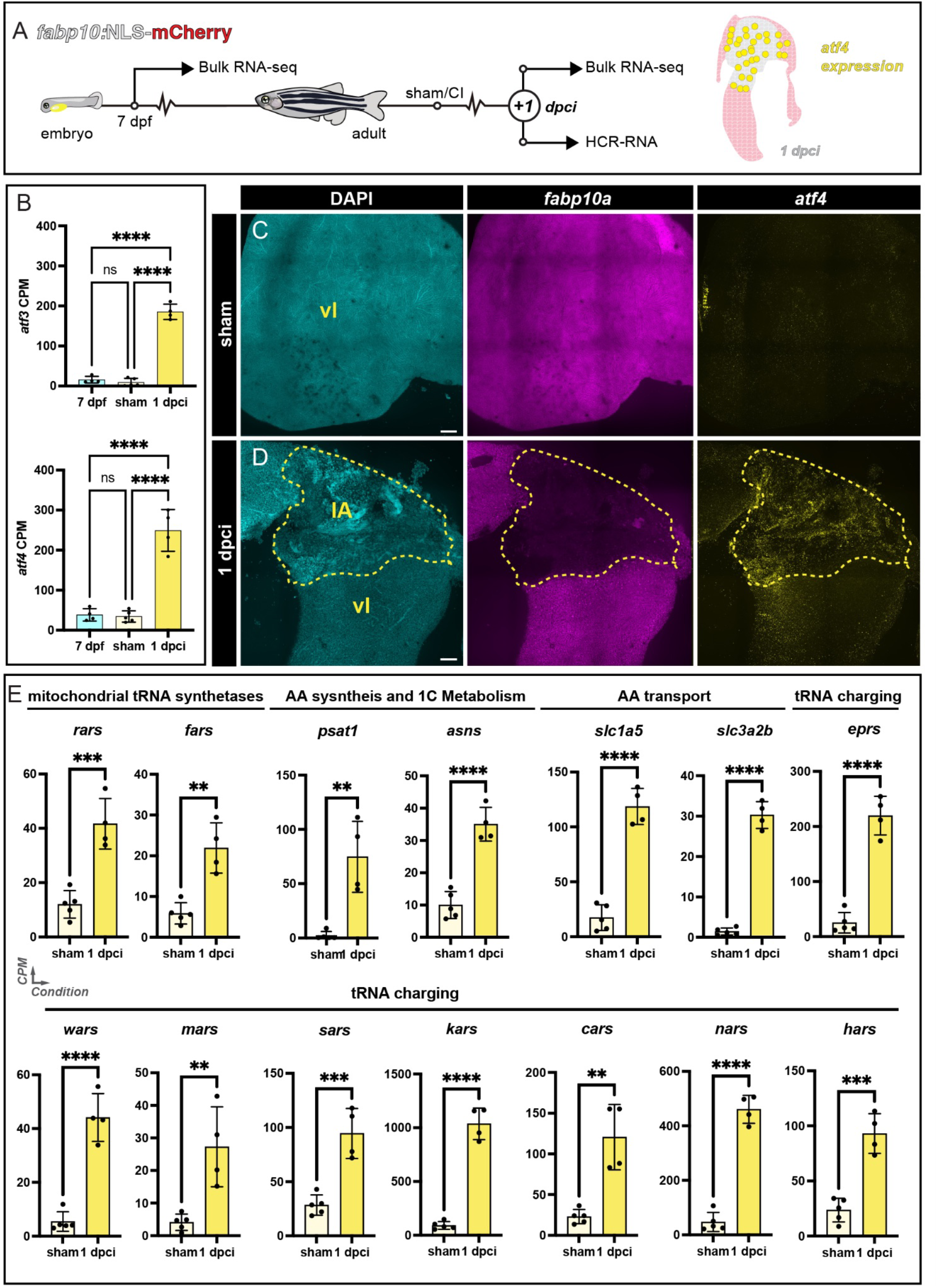
Upregulation of the Integrated stress response (ISR) upon cryoinjury in the zebrafish. A) Graphical illustration of the collection process for sequencing. B) atf4 and atf3 CPM of Bulk RNA-seq livers during developmental (7 dpf), sham and 1 dpci samples. C-D) fapb10a and atf4 HCR staining in sham (C) and 1 dpci (D) adult livers. E) Analysis of Atf4 downstream targets upon cryoinjury in Bulk RNA-seq samples of sham and the IA of 1 dpci adult livers. Dashed line: Border zone of the injured area; dpci: days post-cryoinjury; dpf: days-post fertilisation; IA: injured area; vl: ventral lobe. Scale bars: 200 µm.

